# Patronin condensation integrates recognition, protection and organization of microtubule minus ends

**DOI:** 10.64898/2026.10.03.756418

**Authors:** Boxin Zhang, Junliang Yang, Shuhua Yu, Kun Zhou, Li He, Liangqian Huang, Xin Liang, Heng Wang, Jiong Chen

## Abstract

Microtubule minus ends are central to non-centrosomal microtubule organization, yet how their recognition, protection and molecular organization are integrated remains unclear. Here we show that *Drosophila* Patronin forms dynamic condensates at microtubule minus ends through a prion-like domain (PrLD). PrLD-mediated condensation cooperates with the CKK domain and a second CKK-independent recognition region to convert weak or broadly distributed microtubule-binding activities into persistent minus-end recognition and tracking. Replacing the PrLD with a heterologous phase-separating HNRNPA1 IDR restores minus-end targeting, indicating that condensation capacity rather than PrLD primary sequence is critical. Patronin condensates further form a protective cap that resists minus-end depolymerization and recruit the cytoskeletal regulators Sas-4, Katanin and Ena. Disrupting PrLD multivalency impairs Patronin localization at multiple developmental sites and causes early organismal lethality. Thus, Patronin condensation creates a physiologically essential microtubule minus-end compartment coupling end recognition, stabilization and cytoskeletal organization.

## Introduction

Microtubules are essential cytoskeletal polymers that organize intracellular architecture, transport, cell polarity and division ^1, 2^. Their functions depend critically on the distinct properties of their two ends. The microtubule plus end is highly dynamic, undergoing alternating phases of growth and shrinkage through dynamic instability. Over several decades, extensive work has identified a large network of plus-end-tracking proteins (+TIPs), including EB proteins, CLIP-170 and numerous associated factors, that recognize growing plus ends and regulate their dynamics ^3^. The biochemical interactions, structural determinants and dynamic behaviors underlying plus-end recognition and tracking are now understood in considerable detail. More recently, multivalent interactions and phase separation within +TIP networks have emerged as additional mechanisms that promote selective accumulation at growing microtubule ends and regulate plus-end dynamics ^4–7^.

By contrast, the molecular mechanisms that organize and regulate microtubule minus ends remain much less understood. Whereas plus ends are typically highly dynamic, minus ends in cells are often remarkably stable, displaying little detectable growth or shrinkage. *In vitro*, however, minus ends undergo dynamic polymerization and depolymerization, indicating that there are regulatory proteins in cells to actively keep their stability *in vivo*. Disruption of minus-end stability can strongly alter overall microtubule behavior and increase susceptibility to catastrophe ^8^. In many cells, minus ends are generated and anchored at centrosomal microtubule-organizing centers (MTOCs), where gamma tubulin ring complex (γ-TuRC) promotes nucleation and contributes to minus-end stabilization ^9, 10^. However, many microtubules are not attached to centrosomes, and centrosomal activity is frequently attenuated during differentiation. Such non-centrosomal microtubule arrays therefore require alternative mechanisms to recognize, organize and stabilize their minus ends ^1, 11, 12^.

The CAMSAP/Patronin family has emerged as a central regulator of non-centrosomal microtubule minus ends. CAMSAP/Patronin proteins function as minus-end-targeting proteins (-TIPs) that selectively associate with microtubule minus ends and protect them from depolymerization ^1, 13–19^. A seminal study by Goodwin and Vale demonstrated that depletion of Patronin in *Drosophila* S2 cells causes normally stable minus ends to depolymerize through the action of the Kinesin-13 family depolymerase Klp10A, generating treadmilling microtubules. Purified Patronin selectively bound minus ends and protected them from Kinesin-13-mediated depolymerization *in vitro*, leading to the proposal that Patronin acts as a molecular “cap” that stabilizes the microtubule minus end ^15^. However, the physical nature of this cap remained unresolved.

The same study raised additional mechanistic questions. Although full-length Patronin binds minus ends with high selectivity, its isolated CKK domain, a C-terminal microtubule-binding domain, distributes broadly along the microtubule lattice, indicating that CKK alone cannot account for selective minus-end recognition. Subsequent mammalian studies of CAMSAP1-3 showed that the highly conserved CKK domains bind to minus ends *in vitro*^16, 17^. Moreover, structural work on CAMSAP1 showed that the CKK domain preferentially recognize structural features associated with curved protofilaments near microtubule ends, providing one possible molecular basis for minus-end recognition ^20, 21^. However, when expressed alone in cells, CKK domains of CAMSAP/Patronin proteins generally decorate the microtubule lattice rather than selectively accumulating at minus ends ^15, 22, 23^. Moreover, biochemical studies of *Drosophila* Patronin suggested that a central non-CKK region can itself confer minus-end binding *in vitro* ^16^. Thus, how broadly distributed microtubule-binding activities of CKK domains are converted into robust and persistent minus-end recognition and tracking in cells remains unclear.

Goodwin and Vale also proposed that Patronin might have a broader organizational role at microtubule minus ends. Patronin localized to microtubule-nucleating foci in S2 cells, and overexpressed Patronin or its central coiled-coil region recruited the centrosome-associated proteins Sas-4 and SAK to Patronin-containing structures. These findings suggested that Patronin may function as a scaffold for assembling additional proteins at microtubule nucleation or minus-end sites, but the molecular basis of this scaffolding activity has remained unknown. Thus, more than fifteen years after Patronin was identified as a minus-end regulator, three closely linked questions remain unresolved: how Patronin selectively recognizes and tracks microtubule minus ends, what constitutes the Patronin “cap” that protects these ends from depolymerization, and how Patronin functions as a scaffold to organize additional cytoskeletal regulators at minus ends.

These mechanistic questions have direct physiological relevance. For example, during *Drosophila* oogenesis, the spectraplakin Shot recruits Patronin to the anterior oocyte cortex, where Patronin/Shot foci organize a non-centrosomal microtubule array that establishes anterior-posterior polarity ^24^. Moreover, Patronin also localizes to the apical cortex of follicle cells and act together with Shot to polarize microtubule along apical-basal axis ^24, 25^. Lastly, Patronin’s asymmetric enrichment within the germline cyst contributes to specification of the future oocyte ^26^. How the molecular mechanisms that control Patronin’s minus-end targeting, capping and scaffolding are translated into these stable and spatially restricted localization patterns *in vivo*, and whether the same mechanisms are required for the organismal development, remain unknown.

Liquid-liquid phase separation (LLPS), driven by weak multivalent intermolecular interactions, provides a mechanism by which proteins can assemble into concentrated, dynamic compartments without membranes ^27, 28^. Increasing evidence indicates that phase separation can regulate the microtubule cytoskeleton. Condensation of numerous microtubule-associated proteins has been implicated in microtubule nucleation, branching and regulation of microtubule dynamics ^29–31^. Particularly relevant are recent studies showing that multivalent interactions among +TIPs promote their selective accumulation at growing plus ends, suggesting that condensation can amplify weak molecular preferences into spatially restricted end-associated assemblies ^4–7^. Recent studies further suggest that condensation may be a conserved property of the CAMSAP/Patronin family. Mammalian CAMSAP2 can undergo phase separation with tubulin *in vitro* and promote γ-TuRC-independent microtubule nucleation ^32^. More recently, the *C. elegans* Patronin homolog PTRN-1 was shown to form dynamic condensates in intestinal epithelial cells during recovery from pore-forming toxin injury, but it is not known which region or domain of PTRN-1 is required for condensation and whether PTRN-1 condensates bind microtubule minus-ends ^33^. Therefore, what molecular mechanism drives phase separation of CAMSAP/Patronins, and whether and how phase separation contributes directly to minus-end recognition, tracking, protection, and scaffolding remain unknown.

Here, we show that *Drosophila* Patronin undergoes phase separation *in vivo* and *in vitro* and that this property provides a unifying mechanism for microtubule minus-end organization. We identify a previously uncharacterized prion-like domain (PrLD) that supplies strong multivalent interactions and drives Patronin condensation. PrLD-mediated condensation cooperates with the CKK domain and with a second, CKK-independent minus-end recognition region to convert weak or broadly distributed microtubule-binding activities into robust minus-end recognition and dynamic tracking. The resulting Patronin assembly forms a functional protective cap that stabilizes minus ends against depolymerization and simultaneously acts as a molecular scaffold that enriches additional cytoskeletal regulators, including Sas-4, Katanin and Ena/VASP. Importantly, disrupting PrLD-mediated condensation impairs Patronin enrichment at multiple developmental sites, including the Shot-defined anterior oocyte cortex, the follicle-cell apical cortex and the future oocyte within the germline cyst, and a condensation-deficient Patronin knock-in is homozygous lethal. Together, our findings show that Patronin condensation creates a physiologically essential and multifunctional microtubule minus-end compartment that couples end recognition and tracking with protective capping, cytoskeletal organization and developmental localization.

## Results

### Patronin forms dynamic condensates at microtubule minus ends

To examine Patronin behavior *in vivo*, we first expressed GFP-Patronin in *Drosophila* egg chambers. GFP-Patronin formed puncta that frequently fused (Fig. 1A, B) and also decorated stable microtubules, sometimes coating extended lattice regions ^34^ (Fig. 1A, C), consistent with previous observations ^34^. Both punctate and microtubule-associated Patronin recovered rapidly after photobleaching (Fig. 1C-E), indicating dynamic molecular exchange and suggesting that Patronin forms condensates capable of wetting microtubules. We next examined Patronin in *Drosophila* S2 cells at interphase, which lack a dominant centrosomal microtubule-organizing center (MTOC) and was used extensively by previous mechanistic studies of Patronin^13, 15, 16, 35–37^. mNeonGreen (mNG)-Patronin was expressed from a copper-inducible metallothionein promoter (Fig. 1F). At 25 μM CuSO₄, mNG-Patronin formed microtubule-associated puncta that frequently fused (Fig. 1G). At 50 μM CuSO₄, Patronin formed both puncta and elongated structures; live imaging showed puncta fusing and partially wetting the lattice (Fig. 1H, I). These structures often resolved into bead-like droplets, consistent with Rayleigh–Plateau-like instability of a liquid coating on a filament (Fig. 1I, at 0”) ^38^. At higher expression, Patronin extensively wetted microtubules (Extended Data Fig. 1A, B), while microtubule-associated Patronin showed rapid FRAP recovery (Fig. 1J, K).

**Fig. 1.**
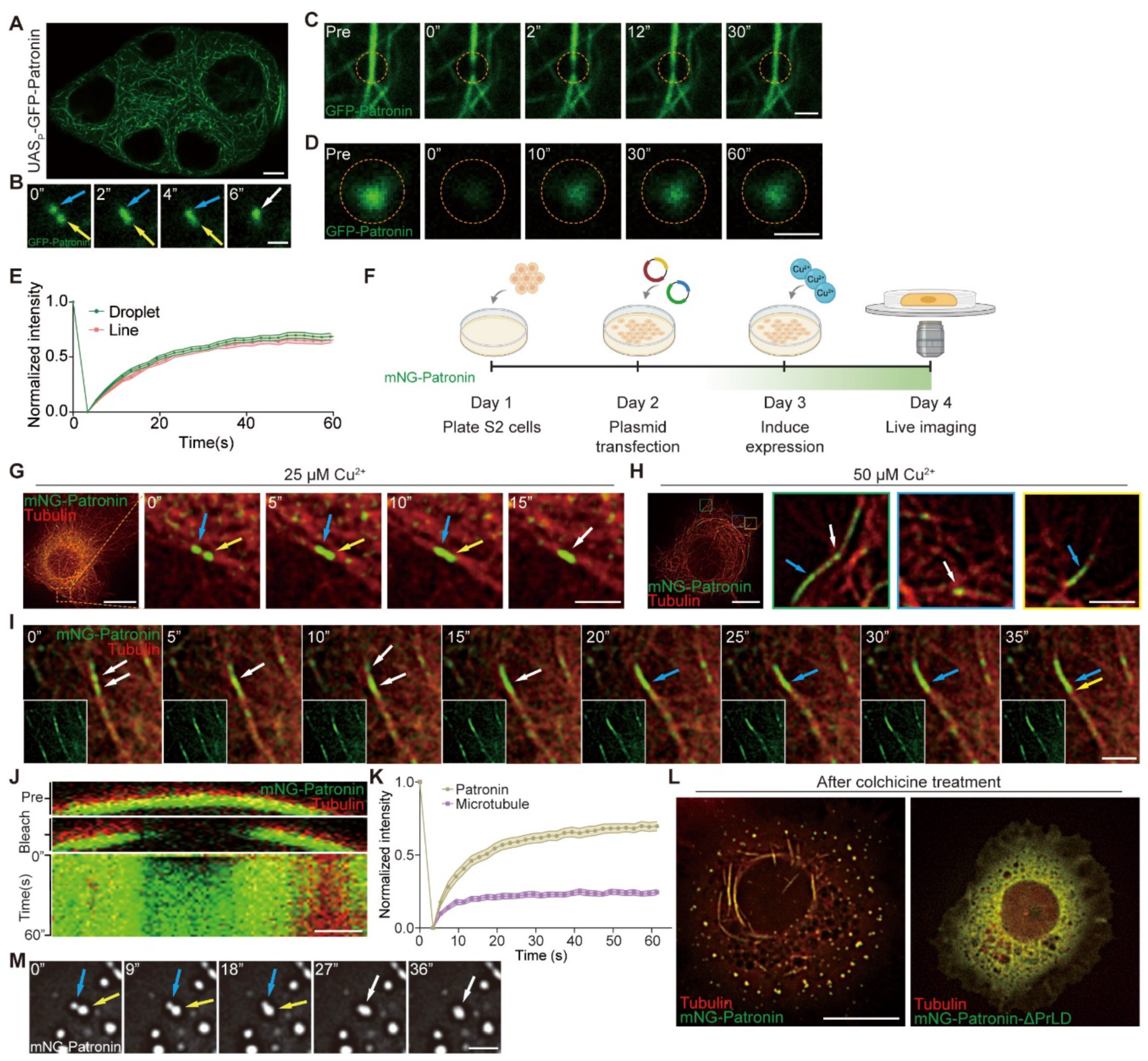
Patronin forms dynamic condensates in egg chambers and S2 cells (A) The distribution pattern of GFP-Patronin in *Drosophila* stage 7 egg chambers. *UASp-GFP-Patronin* was expressed in the germline by the *Mat-Gal4* driver. Scale bar, 10 μm. (B) Live imaging of GFP-Patronin droplets undergoing fusion in egg chamber from 0 to 6 seconds. 6” represents 6 seconds. Scale bar, 1 μm. (C-E) FRAP of a GFP-Patronin enriched line (C) and a GFP-Patronin enriched droplet or punctae (D) in egg chambers, FRAP curves are plotted in (E). Data are presented as mean ± s.e.m. of at least 30 samples per time point for each curve. Scale bars, 2 μm (C), 1 μm (D). (F) Schematic of live imaging in S2 cells, illustrating the copper-inducible expression plasmid strategy used throughout this study. (G) Live imaging of mNG-Patronin droplet fusion on microtubules in S2 cells. mNG-Patronin expression is induced with 25 µM Cu^2+^, while mCherry-αTub84B was expressed by actin promoter to label microtubules throughout this study. The low magnification view is on the left and a boxed region within it is enlarged and shown to its right as a series of time-lapsed images. Scale bars,10 μm (left) and 2 μm (right). (H) Live imaging of mNG-Patronin on microtubules in S2 cells induced with 50 μM Cu^2+^. Unless otherwise stated, all other experiments in this study involving S2 cells used 50 μM Cu^2+^ for induced expression. Boxed regions in blue, green, and yellow (left panel) are enlarged and shown on the right, each representing a distinct pattern. White arrows indicate puncta, and blue arrows point to rod-like or elongated structures. Scale bars,10 μm (left) and 2 μm (right). (I) Time-lapse imaging of mNG-Patronin on microtubules in S2 cells. White arrows indicate fusion of punctate Patronin from 0 to 15 seconds, while blue and yellow arrows indicate Patronin wetting microtubules from 20 to 35 seconds. Yellow arrow indicates the extent of wetting. Scale bar, 2 μm. (J, K) High mNG-Patronin expression was induced with 200 µM Cu^2+^, FRAP was used to quantify the dynamics of mNG-Patronin on microtubules. Kymographs show rapid recovery of Patronin fluorescence over time, whereas microtubule fluorescence exhibits little recovery. Data are presented as mean ± s.e.m. from at least 30 samples per time point. Scale bar, 5 μm. (L) Distribution of Patronin and Patronin-ΔPrLD in S2 cells following microtubule depolymerization with 200 μM colchicine for 6 h. Scale bar, 10 μm. (M) Upon microtubule depolymerization, Patronin forms droplets capable of fusing with one another in the cytoplasm of S2 cells. Scale bar, 2 μm.

Patronin condensation did not require microtubule array. After colchicine-induced microtubule depolymerization, Patronin formed fusible cytoplasmic droplets (Fig. 1L, left panel, and 1M), demonstrating an intrinsic capacity to condense independently of the lattice. We confirmed this property with purified protein. MBP-Patronin-GFP purified from Sf9 cells remained soluble before MBP removal (Fig. 2A and Extended Data Fig. 1C), but after cleavage Patronin-GFP spontaneously formed condensates at physiological salt concentrations without molecular crowding agents (Fig. 2B). Condensation was detectable at 10 nM and increased with protein concentration (Fig. 2B, C). The condensates underwent fusion and rapid FRAP recovery (Fig. 2D, E), demonstrating intrinsic dynamic condensation *in vitro*.

**Fig. 2.**
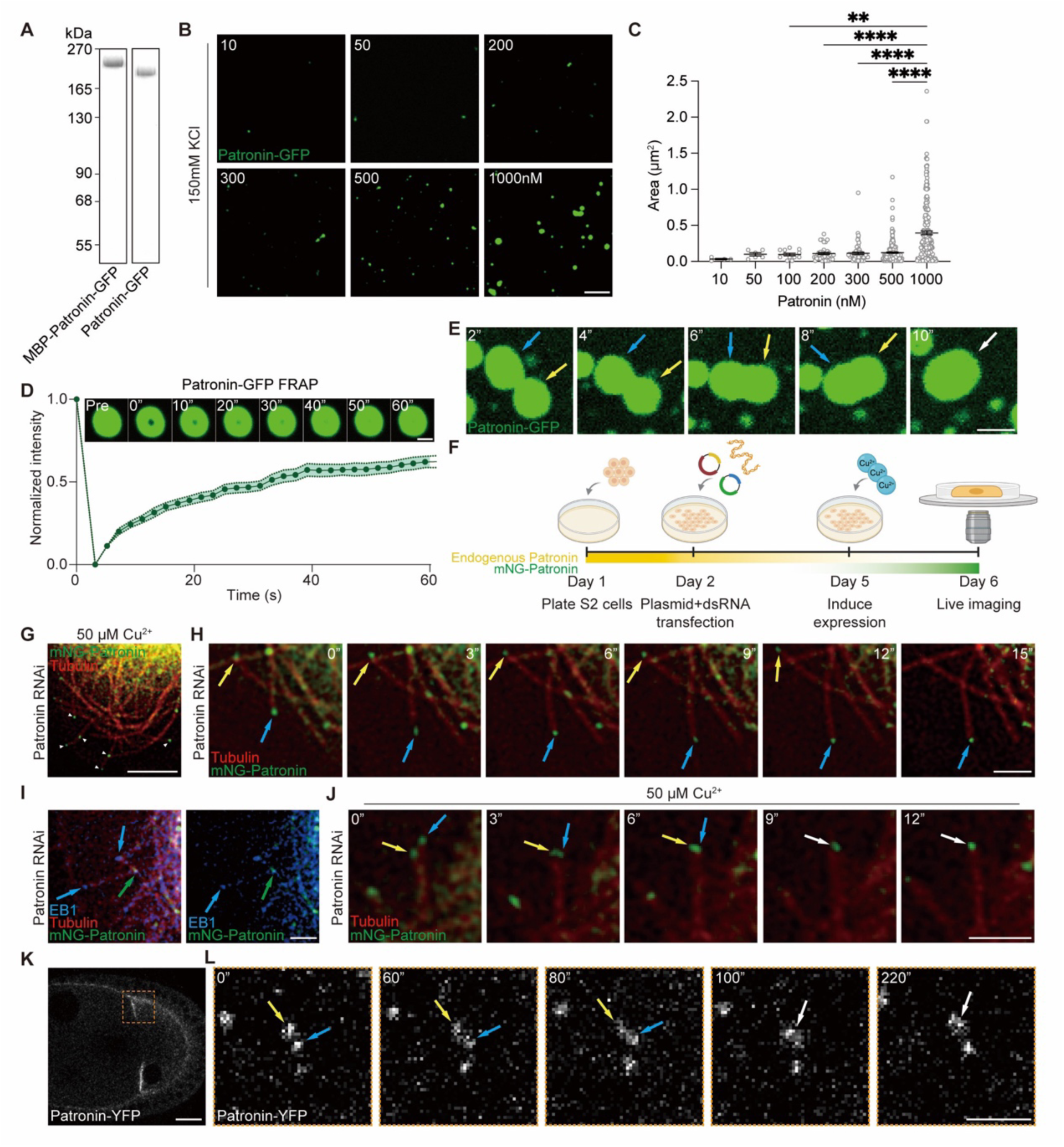
Patronin undergoes phase separation at the microtubule minus ends. (A) Coomassie Brilliant Blue staining of purified MBP-Patronin-GFP and Patronin-GFP (after MBP cleavage). (B) Representative fluorescence images of Patronin-GFP phase separating at different protein concentrations *in vitro* in a buffer containing 150 mM KCl. Scale bar, 5 μm. (C) Quantification of cross-sectional area of Patronin droplets in (B). This and all other multiple comparisons in this study were analyzed by one-way ANOVA. Data are presented as means ± SEM, with statistical significance set as follows: *\*P < 0.05; **P < 0.01; ***P < 0.001; ****P < 0.0001*. (D) FRAP analysis of Patronin-GFP droplets (1000 nM) *in vitro*. Top: time-lapse images from a FRAP experiment. Bottom: fluorescence recovery curves over time after photobleaching within droplets. Data are presented as mean ± s.e.m. for each time point; n = 30. Scale bar, 1 μm. (E) Representative images of Patronin-GFP droplet fusion. Scale bar, 1 μm. (F) Schematic of *Patronin RNAi* in S2 cells, illustrating the strategy used to knock down endogenous Patronin in this study. (G) Representative images of mNG-Patronin expressed in S2 cells depleted of endogenous Patronin. White arrowheads indicate Patronin condensates at microtubule minus ends. Scale bar, 5 μm. (H) Time-lapse imaging of mNG-Patronin in S2 cells depleted of endogenous Patronin. Blue and yellow arrows indicate Patronin condensates tracking microtubule minus ends, which are dynamic and moving. Scale bar, 2 μm. (I) Representative images of mNG-Patronin and EB1-BFP (blue) in S2 cells depleted of endogenous Patronin. Blue arrows indicate EB1-labeled microtubule plus ends, and green arrows indicate Patronin-labeled minus ends. Scale bar, 2 μm. (J) Live imaging of mNG-Patronin condensate fusion in S2 cells depleted of endogenous Patronin. Blue and yellow arrows point to two Patronin condensates before or during fusion, and white arrow indicates Patronin condensate after fusion. Scale bar, 2 μm. (K, L) Live imaging of Patronin-YFP condensates fusing in stage 9 egg chambers. Yellow boxed region in (K) is magnified in (L). Blue and yellow arrows point to two Patronin condensates before and during fusion, and white arrow indicates Patronin condensate after fusion. Scale bars, 20 μm (left) and 3 μm (right).

We then asked whether condensates form at minus ends at physiological or near-physiological Patronin levels. After depleting endogenous Patronin, low-level mNG-Patronin induced with 50 μM CuSO₄ accumulated selectively at one microtubule end (Fig. 2F-H). EB1 marked the opposite, plus end, confirming minus-end localization (Fig. 2I). Minus-end-associated Patronin puncta also fused (Fig. 2J), indicating condensate-like behavior at the minus end. Endogenous Patronin behaved similarly *in vivo*. In stage 9 oocytes from a Patronin-YFP knock-in line, Patronin-YFP formed puncta at the anterior cortex ^24^, where microtubule minus ends are enriched ^24^ (Fig. 2K), and neighboring puncta fused during live imaging (Fig. 2L). Thus, Patronin forms dynamic condensates in purified systems and cells, and these condensates selectively accumulate at microtubule minus ends at physiological or near-physiological levels.

### A prion-like domain provides the multivalent interactions required for Patronin condensation

To define the molecular basis of Patronin condensation, we analyzed its sequence and identified an extensive intrinsically disordered region (IDR) containing a previously uncharacterized ∼300-amino-acid prion-like domain (PrLD) (Fig. 3A-C). PrLDs commonly contain multivalent interaction motifs that promote biomolecular condensation ^39^. Deleting the PrLD strongly impaired Patronin condensation. Unlike wild-type Patronin, which formed punctate and elongated assemblies on microtubules, Patronin-ΔPrLD distributed uniformly along the lattice in S2 cells (Fig. 3D) and failed to form cytoplasmic condensates after colchicine-induced microtubule depolymerization (Fig. 1L, right panel). Thus, the PrLD is required for both microtubule-associated and microtubule-independent condensation.

**Fig. 3.**
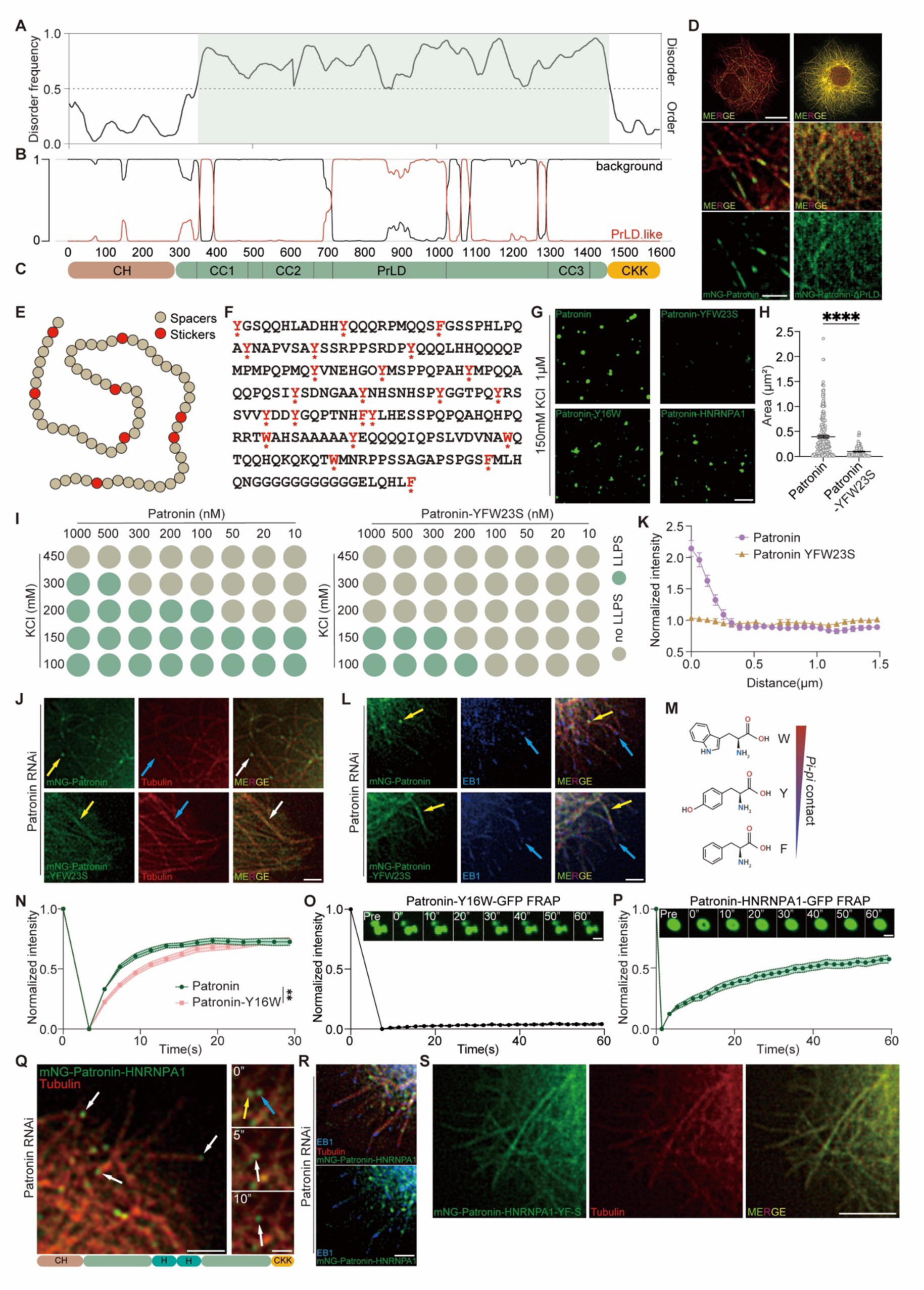
A prion-like domain provides the multivalent interactions and drives Patronin’s phase separation. (A-C) Schematic of *Drosophila* Patronin domain architecture (C) and sequence features. (A) The line at 0.5 (y axis) is the cut-off for disorder (>0.5) and order (<0.5) predictions, which is predicted by VL3, a prediction program for disordered dispositions. (B) Patronin contains a prion-like domain that is predicted by the PLAAC prion prediction program. (D) Representative images of S2 cells expressing Patronin or Patronin-ΔPrLD. Middle and bottom panels are magnified views. Microtubules are labeled by mCherry-αTub84B (red). Scale bars,10 μm (upper panels), 2 μm (lower panels). (E) Schematic representation of the stickers-and-spacers model. (F) Amino acid sequence of the Patronin PrLD. The 23 aromatic residues that act as stickers are indicated by red asterisks. (G) *In vitro* LLPS (liquid-liquid phase separation) behavior of Patronin, Patronin mutants and Patronin–HNRNPA1 chimeras at 1 μM in 150 mM KCl. Scale bar, 5 μm. (H) Comparison of droplet areas between Patronin-GFP (n=247) and Patronin-YFW23S-GFP (n=117) at 1000 nM in 150 mM KCl. For this and other comparisons between two groups in this study, unpaired two-tailed Student’s t-tests were used. Data are presented as means ± SEM, with statistical significance set as follows: *\*P < 0.05; **P < 0.01; ***P < 0.001; ****P < 0.0001*. (I) Summary of *in vitro* LLPS behavior of Patronin and Patronin-YFW23S. Green circles indicate phase separation; brown circles indicate no phase separation. (J, K) Representative images of S2 cells depleted of endogenous Patronin and expressing mNG-Patronin or mNG-Patronin-YFW23S. (K) shows quantification of Patronin (n = 15) and Patronin-YFW23S (n = 13) fluorescence along a 1.5-μm segment from individual microtubule ends. Data are presented as mean ± s.e.m. at each position. Scale bar, 2 μm. (L) Representative images of S2 cells depleted of endogenous Patronin and co-expressing mNG-Patronin or mNG-Patronin-YFW23S with EB1-BFP. Yellow arrows, Patronin or Patronin-YFW23S; blue arrows, EB1. Scale bar, 2 μm. (M-O) Tyrosine-to-Tryptophan substitutions increased pi-pi interactions (M) within the PrLD, resulting in altered condensate morphology (O, upper panel) and reduced condensate dynamics in both S2 cells (N) and *in vitro* (O). FRAP data are presented as mean ± s.e.m. at each time point; n = 30. 1 μM of Patronin-Y16W-GFP used for FRAP. Scale bar, 1 μm. (P) FRAP analysis of Patronin-HNRNPA1-GFP droplets (1 μM) in vitro. Top: time-lapse images from a FRAP experiment. Bottom: fluorescence recovery curves over time after photobleaching within droplets. Data are presented as mean ± s.e.m. for each time point; n = 30. Scale bar, 1 μm. (Q) Representative images of S2 cells depleted of endogenous Patronin and expressing mNG-Patronin-HNRNPA1. Right, magnified views showing time-lapse imaging of Patronin-HNRNPA1 condensates over 10 seconds. Yellow and blue arrows indicate condensates before fusion; white arrows indicate fused condensates. Bottom, schematic of the Patronin-HNRNPA1 chimera; H indicates HNRNPA1 IDR. Note that there are two tandem copies of HNRNPA1 IDR, whose combined size of 268 amino acids is comparable to that of PrLD (312 amino acids). Scale bars, 2 μm (left panel), and 1 μm (right panels). (R) Representative images of S2 cells depleted of endogenous Patronin and co-expressing mNG-Patronin-HNRNPA1 with EB1. Scale bar, 2 μm. (S) Representative images of S2 cells expressing mNG-Patronin-HNRNPA1-YF-S. Scale bar, 5 μm.

The PrLD contains numerous aromatic residues interspersed among polar residues, consistent with a sticker-and-spacer architecture (Fig. 3E, F)^39^. Replacing all 23 aromatic residues with serine (Patronin-YFW23S) markedly reduced condensation of purified Patronin and increased the concentration required for condensate formation (Fig. 3G-I and Extended Data Fig. 2B). In S2 cells, Patronin-YFW23S likewise failed to form punctate or rod-like structures and instead distributed uniformly along microtubules (Extended Data Fig. 2C). Disrupting PrLD interactions also abolished minus-end enrichment: after endogenous Patronin depletion, Patronin-YFW23S decorated the microtubule lattice rather than accumulating at minus ends (Fig. 3J-L). Mutating aromatic residues in either half of the PrLD produced the same phenotype (Extended Data Fig. 2D), linking PrLD multivalency and condensation to minus-end targeting. Conversely, strengthening pi-pi interactions within PrLD by replacing tyrosines with tryptophans (Fig. 3M), generated Patronin-Y16W, which showed slower FRAP recovery in S2 cells (Fig. 3M, N) and formed less dynamic, solid-like assemblies *in vitro* (Fig. 3G, O). Thus, aromatic interaction strength tunes Patronin condensate material properties. To test whether minus-end targeting requires the Patronin PrLD sequence itself or simply multivalent condensation, we replaced the PrLD with two tandem copies of the phase-separating HNRNPA1 IDR (Extended Data Fig. 2E), which was previously shown to be a prion-like domain and contain aromatic residues-dependent sticker-and-spacer architecture that promotes LLPS ^39, 40^. Patronin-HNRNPA1 formed dynamic condensates *in vitro* (Fig. 3G, P) and fusible minus-end-enriched condensates in Patronin-depleted S2 cells (Fig. 3Q, R). Mutating aromatic residues in the HNRNPA1 IDR abolished minus-end localization and redistributed the chimera uniformly along microtubules (Fig. 3S). Together, these data identify the PrLD as a major driver of Patronin condensation and show that condensation capacity, rather than the PrLD primary sequence, is critical for robust minus-end enrichment.

### PrLD-mediated condensation converts CKK binding into minus-end recognition and tracking

We next asked whether condensation itself recognizes the minus end or instead enables another Patronin domain to do so. The isolated PrLD formed condensates *in vitro* in the presence of a crowding agent, whereas aromatic-residue mutation abolished condensation (Fig. 4A-C). In S2 cells, blue light induced fusible Opto-PrLD condensates but not Opto-PrLD-YFW23S condensates (Fig. 4D-G). Neither isolated PrLD nor Opto-PrLD associated detectably with microtubules (Fig. 4E, G, H and Extended Data Fig. 3), showing that condensation alone is insufficient for minus-end recognition. The CKK domain provided the complementary activity. CKK bound microtubules in S2 cells but distributed broadly along the lattice and diffusely in the cytoplasm rather than accumulating at minus ends (Fig. 4A, I), consistent with previous study ^15^. Purified CKK also failed to condense *in vitro*, even with a crowding agent (Fig. 4J). Thus, CKK binds microtubules in S2 cells but lacks robust minus-end specificity on its own.

**Fig. 4.**
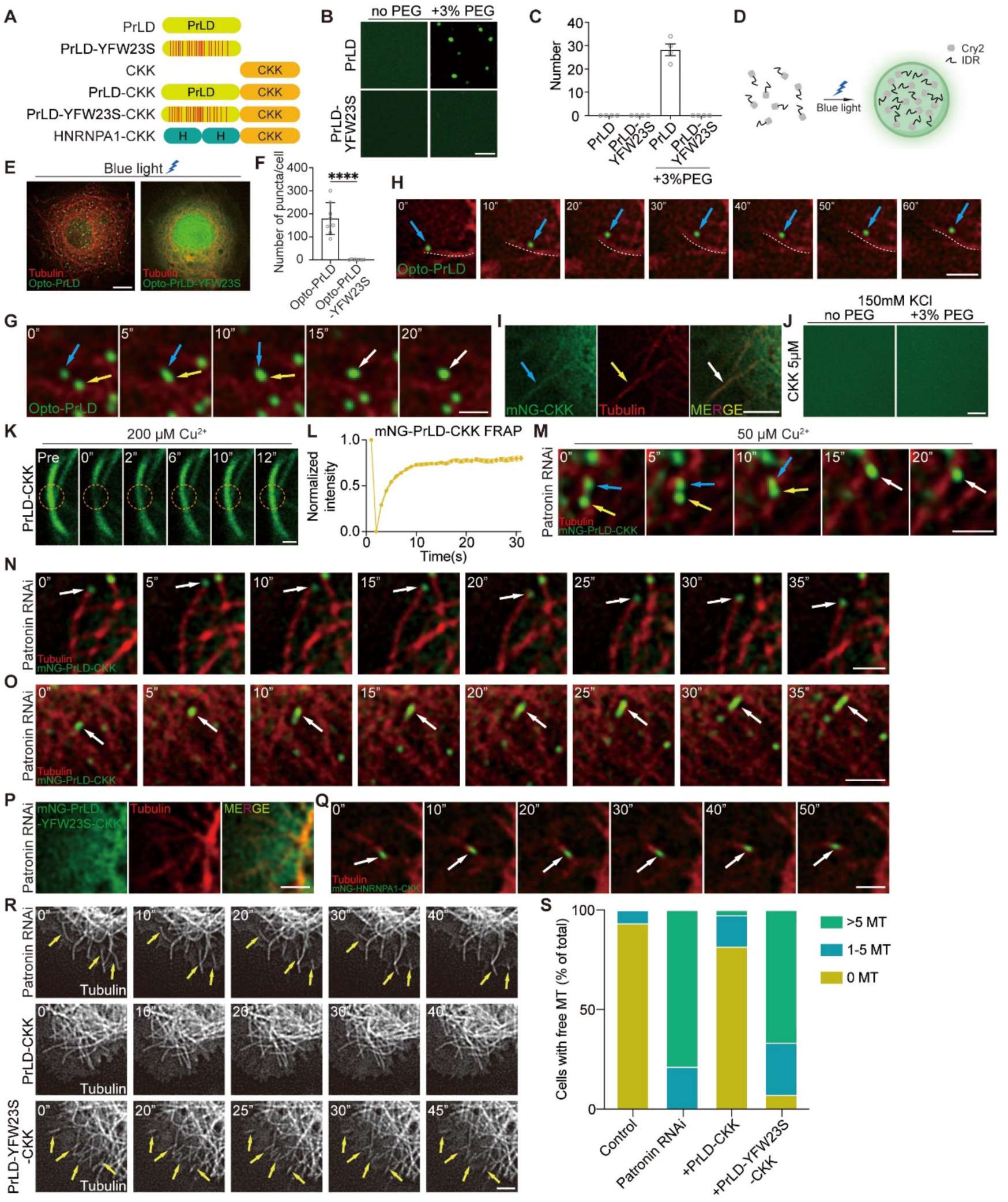
PrLD-mediated condensation cooperates with CKK in minus-end targeting. (A) Schematic of Patronin truncations and mutants. (B, C) Summary of *in vitro* LLPS behavior of PrLD and PrLD mutants at 2 μM in 150 mM KCl with or without crowding agent. (C) is a quantification of (B). Scale bar, 3 μm. (D) Schematic of optogenetic induction of droplets in S2 cells. (E, F) Representative images of S2 cells expressing opto-PrLD or opto-PrLD-YFW23S under blue-light illumination for 30 s (E), and quantified in (F). Scale bar, 5 μm. (G, H) Live imaging showing blue light-induced Opto-PrLD condensates. Yellow and blue arrows in (G) indicate droplets before and during fusion, white arrows indicate droplet after fusion. Dashed line in (H) indicates a microtubule, blue arrows indicate an Opto-PrLD droplet. Notably, Opto-PrLD droplets is adjacent to a microtubule, but did not stably bind to it nor track its minus end. Scale bars, 1 μm (G), 2 μm (H). (I) Representative images of S2 cells expressing mNG-CKK. Scale bar, 2 μm. (J) No *in vitro* LLPS behavior is detected for CKK at 5 μM in 150 mM KCl with or without a crowding agent. Scale bar, 10 μm. (K, L) FRAP analysis of mNG-PrLD-CKK dynamics on microtubules in S2 cells after induced expression with 200 μM Cu^2+^. Scale bar, 1 μm. (M) Live imaging of fusion of mNG-PrLD-CKK condensates in S2 cells depleted of endogenous Patronin. Scale bar, 1 μm. (N, O) Live imaging of mNG-PrLD-CKK in S2 cells depleted of endogenous Patronin. White arrows in (N) indicate PrLD-CKK condensates tracking the moving microtubule minus ends. White arrows in (O) indicate PrLD-CKK condensates wetting microtubules and forming stretches. Scale bars, 1 μm (N), 2 μm (O). (P, Q) Live imaging of S2 cells depleted of endogenous Patronin shows that mNG-PrLD-YFW23S-CKK lost minus-end binding (P), whereas mNG-HNRNPA1-CKK restored minus-end tracking (Q). White arrows indicate HNRNPA1-CKK condensates that bind to and track microtubule minus ends. Scale bar, 1 μm. (R, S) Time-lapse imaging of Patronin-depleted S2 cells alone (n=33), or co-expressing PrLD-CKK (n=38), or co-expressing PrLD-YFW23S-CKK (n = 42). Yellow arrows in (R) indicate the free microtubules that are dynamic and undergoing treadmilling. Also see Supplementary Movie 1. Scale bar, 2 μm. Quantification of free microtubules per cell is shown in (S). Colored bars represent the percentage of cells observed with the indicated number of free microtubules (n = total number of cells).

Coupling the PrLD and CKK activities was sufficient to restore minus-end targeting. At high expression, PrLD-CKK formed dynamic condensates that wetted microtubules and displayed rapid recovery after photobleaching (Fig. 4A, K, L). After endogenous Patronin depletion and at lower expression, PrLD-CKK formed fusible condensates that selectively accumulated at and dynamically tracked microtubule minus ends (Fig. 4M, N). It also extended a short distance along the lattice from the minus end, forming the characteristic CAMSAP “stretch” ^17^ (Fig. 4O). Thus, neither PrLD-mediated condensation nor CKK-mediated microtubule binding is sufficient alone; together they reconstitute selective minus-end recognition and tracking. This activity required condensation: PrLD-YFW23S-CKK lost minus-end preference (Fig. 4P), whereas replacing the Patronin PrLD with the heterologous phase-separating HNRNPA1 IDR restored selective minus-end localization (Fig. 4Q). These results show that CKK provides microtubule-binding and minus-end-recognition information^17, 20^, while PrLD-mediated condensation amplifies this weak preference into stable, persistent minus-end targeting.

### Patronin condensation generates a functional cap that protects microtubule minus ends

Patronin was previously proposed to form a “cap” that protects microtubule minus ends from Kinesin-13-mediated depolymerization ^15^. We therefore tested whether PrLD-mediated condensation is required for this protective activity. Patronin depletion generated numerous treadmilling microtubules (Fig. 4R, upper panel) ^15^, reflecting plus-end growth coupled to Kinesin-13-mediated depolymerization of unprotected minus ends ^15, 41^. PrLD-CKK efficiently suppressed treadmilling and restored stable microtubule behavior (Fig. 4R, S), whereas condensation-deficient PrLD-YFW23S-CKK failed to rescue despite retaining CKK. Thus, CKK-mediated microtubule binding alone is insufficient for efficient minus-end stabilization; PrLD-mediated condensation converts CKK-dependent binding into a protective activity that resists depolymerization. Together with the localization and wetting data, these results provide a physical interpretation of the Patronin cap. Rather than acting only as a small stoichiometric complex on terminal tubulin subunits, Patronin can assemble a condensate-based functional cap encompassing the minus end and adjacent stretch. This mesoscale compartment provides persistent minus-end occupancy and protection, linking minus-end recognition, tracking and capping in a single mechanism.

### CC3-mediated condensation enhances CKK function but cannot confer minus-end specificity

Because Patronin-YFW23S retained residual condensation despite strong PrLD disruption (Fig. 3G, H), we asked whether Patronin contains another phase-separation module. The CC3-CKK region of mammalian CAMSAP2 can condense *in vitro* ^32^. Purified Patronin CC3-CKK likewise formed condensates at physiological salt concentrations without crowding agents, although at substantially higher protein concentrations than full-length Patronin (Fig. 5A-C). These condensates recovered after photobleaching and were salt-sensitive (Fig. 5D and Extended Data Fig. 4A), consistent with dynamic assemblies driven by weak interactions. Truncation analysis localized the major activity to CC3 and the linker region between CC3 and CKK (Fig. 5E): R1, R2 and R3 formed condensates under different *in vitro* conditions, whereas isolated CKK did not (Fig. 4J, 5F-H and Extended Data Fig. 4B). However, removing CKK weakened condensation relative to CC3-CKK (Fig. 5F-H), indicating that CKK cooperatively enhances CC3-driven assembly despite being unable to condense alone. CC3-mediated condensation also enhanced CKK function. CC3-CKK associated with microtubules more efficiently than CKK alone in S2 cells (Fig. 5I, J). At low expression, CKK failed to rescue Patronin-depletion-induced treadmilling, whereas CC3-CKK and condensate-forming R1 and R2 restored normal microtubule behavior (Fig. 5K, L and Extended Data Fig. 4C). High CKK expression partially rescued treadmilling (Extended Data Fig. 4D, E), while R3 condensed but lacked CKK and failed to rescue at either low or high expression (Fig. 5L and Extended Data Fig. 4C, E). Thus, CC3-mediated condensation increases the functional efficiency of CKK, enhancing microtubule binding and stabilization at relatively low protein levels.

**Fig. 5.**
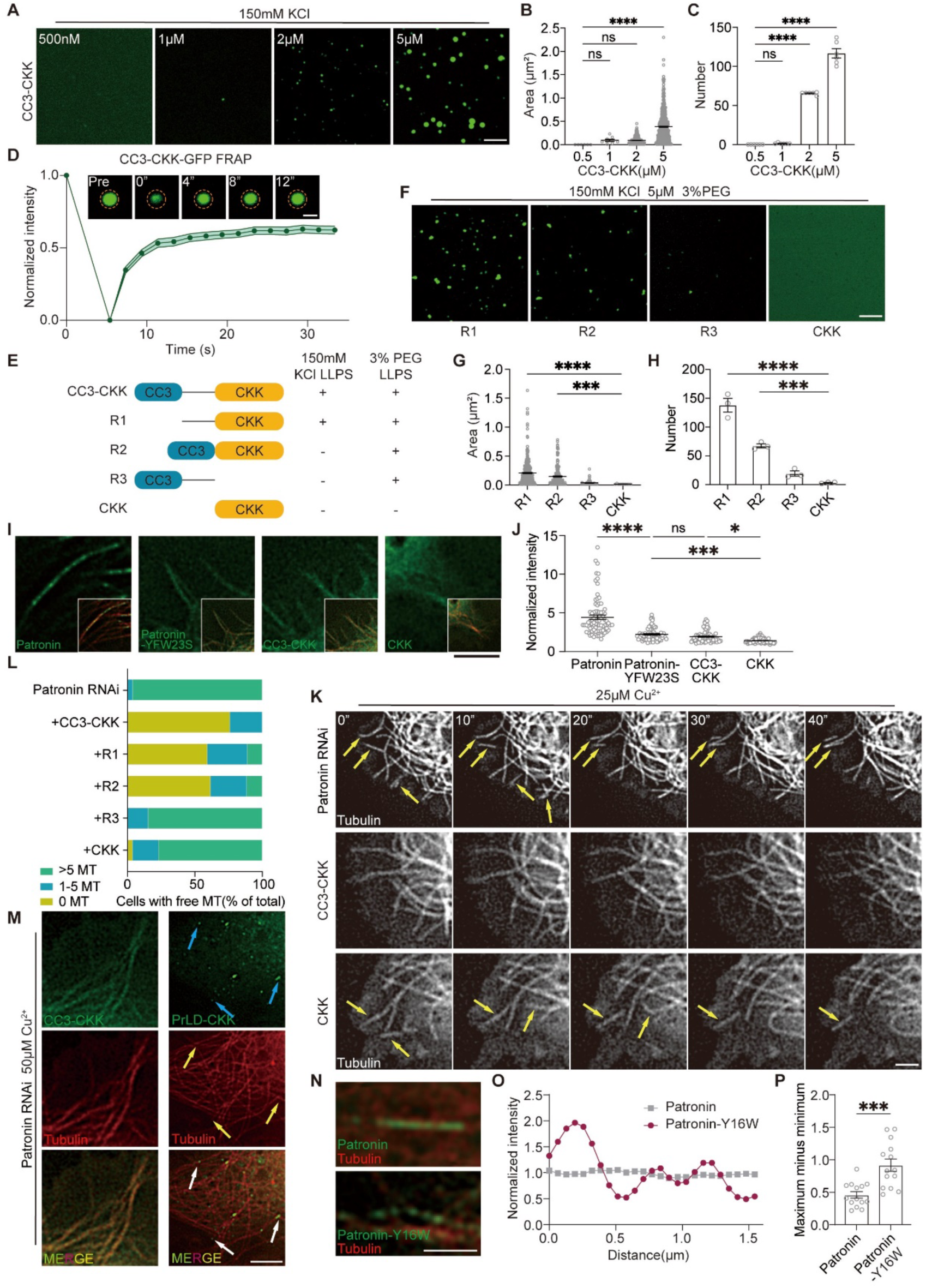
CC3-mediated condensation enhances CKK function. (A-C) Representative fluorescence images of CC3-CKK-GFP (A), phase separating at different protein concentrations *in vitro* in a buffer containing 150 mM KCl. Scale bar, 5 μm. Quantification of CC3-CKK droplet area (B) and number of condensates (C) is performed. (D) FRAP analysis of CC3-CKK-GFP droplets (2 μM) *in vitro*. Top: time-lapse images from a FRAP experiment. Bottom: fluorescence recovery curves over time after photobleaching within droplets. Scale bar, 1 μm. (E-H) Schematic of CC3-CKK truncations and summary of their *in vitro* LLPS behavior (5 μM), under the indicated salt conditions with or without crowding agent (E). −, no LLPS observed; +, LLPS observed. Representative *in vitro* LLPS images of CC3-CKK truncations are shown in (F), and Quantification of CC3-CKK droplet area (G) and number of condensates (H) is shown. Scale bar, 5 μm. (I, J) Representative images of S2 cells expressing Patronin, Patronin-YFW23S, CC3-CKK or CKK (I), under the induction of 100 μM Cu^2+^. Insets (bottom right): microtubules are labeled with mCherry-αTub84B (red). Scale bar, 2 μm. (J) Quantification of the microtubule-to-cytoplasm fluorescence ratio, which indicates how well each Patronin variant bound microtubule. (K, L) (K) Live imaging of S2 cells expressing *Patronin RNAi* alone, *Patronin RNAi* and CC3-CKK, or *Patronin RNAi* and CKK, under induction with 25 μM Cu^2+^. Yellow arrows indicate “free microtubules” that are dynamic and treadmilling. Scale bar, 2 μm. Also see Supplementary Movie 2. (L) Quantification of free microtubules per cell from *Patronin RNAi* alone (n = 26), or expressing *Patronin RNAi* together with CC3-CKK (n = 25), R1 (n = 26), R2 (n = 27), R3 (n = 28), or CKK (n = 26). n, number of cells imaged. (M) Representative images of S2 cells depleted of endogenous Patronin and expressing mNG-CC3-CKK or mNG-PrLD-CKK. Arrows indicate mNG-PrLD-CKK condensates at microtubule minus ends. Scale bar, 5 μm. (N-P) Representative images of S2 cells expressing Patronin or Patronin-Y16W (N). Scale bar, 2 μm. These two images are analyzed in (O), in which Patronin or Patronin-Y16W condensate fluorescence along a 1.5-μm segment of microtubule was measured and plotted. (P) Quantification of the beads-on-the-string morphology of Patronin and Patronin-Y16W condensates. The maximum intensity of a wave (e.g. measured from a 1.5-μm segment of microtubule in O) minus the minimum intensity is used as a measurement for beads-on-the-string morphology. Data are mean ± s.e.m.; n = 14.

CC3-mediated condensation, however, did not confer PrLD-like minus-end specificity. CC3-CKK bound microtubules similarly to Patronin-YFW23S and less efficiently than wild-type Patronin (Fig. 5I, J); after Patronin depletion, it decorated the lattice rather than selectively accumulating at minus ends (Fig. 5M), whereas PrLD-CKK robustly targeted minus ends (Fig. 5M). Thus, weaker CC3-mediated condensation increases CKK occupancy on microtubules, whereas stronger PrLD-mediated condensation converts CKK binding into minus-end specificity. The two modules therefore likely differ in cohesive or interfacial properties. Consistent with this, strengthening PrLD aromatic interactions by tyrosine-to-tryptophan substitution increased Rayleigh-Plateau-like breakup ^38^ of Patronin assemblies into bead-like droplets (Fig. 5N-P). Although this does not directly measure surface tension, it is consistent with stronger cohesion and altered interfacial properties of PrLD-driven condensates. Together, Patronin contains at least two condensation modules with distinct outputs: weak CC3-mediated condensation enhances microtubule binding, whereas stronger PrLD multivalency is required for selective minus-end enrichment.

### A second microtubule-recognition module cooperates with PrLD condensation to track minus ends

Although the results above explain CKK-dependent minus-end targeting, previous work suggested that Patronin also contains a CKK-independent minus-end-binding activity ^16^. We therefore tested whether another Patronin region can provide minus-end recognition. Patronin-ΔCKK was largely diffuse in the cytoplasm and along microtubules and formed only small residual condensates (Fig. 6A), making minus-end specificity difficult to assess because CKK deletion reduced both microtubule association and condensation. We therefore restored condensation optogenetically. Blue light induced fusible Opto-Patronin-ΔCKK condensates (Fig. 6B and Extended Data Fig. 5A) that preferentially associated with one microtubule end (Fig. 6C). EB1 marked the opposite plus end, confirming that these condensates recognized minus ends despite lacking CKK (Fig. 6D).

**Fig. 6.**
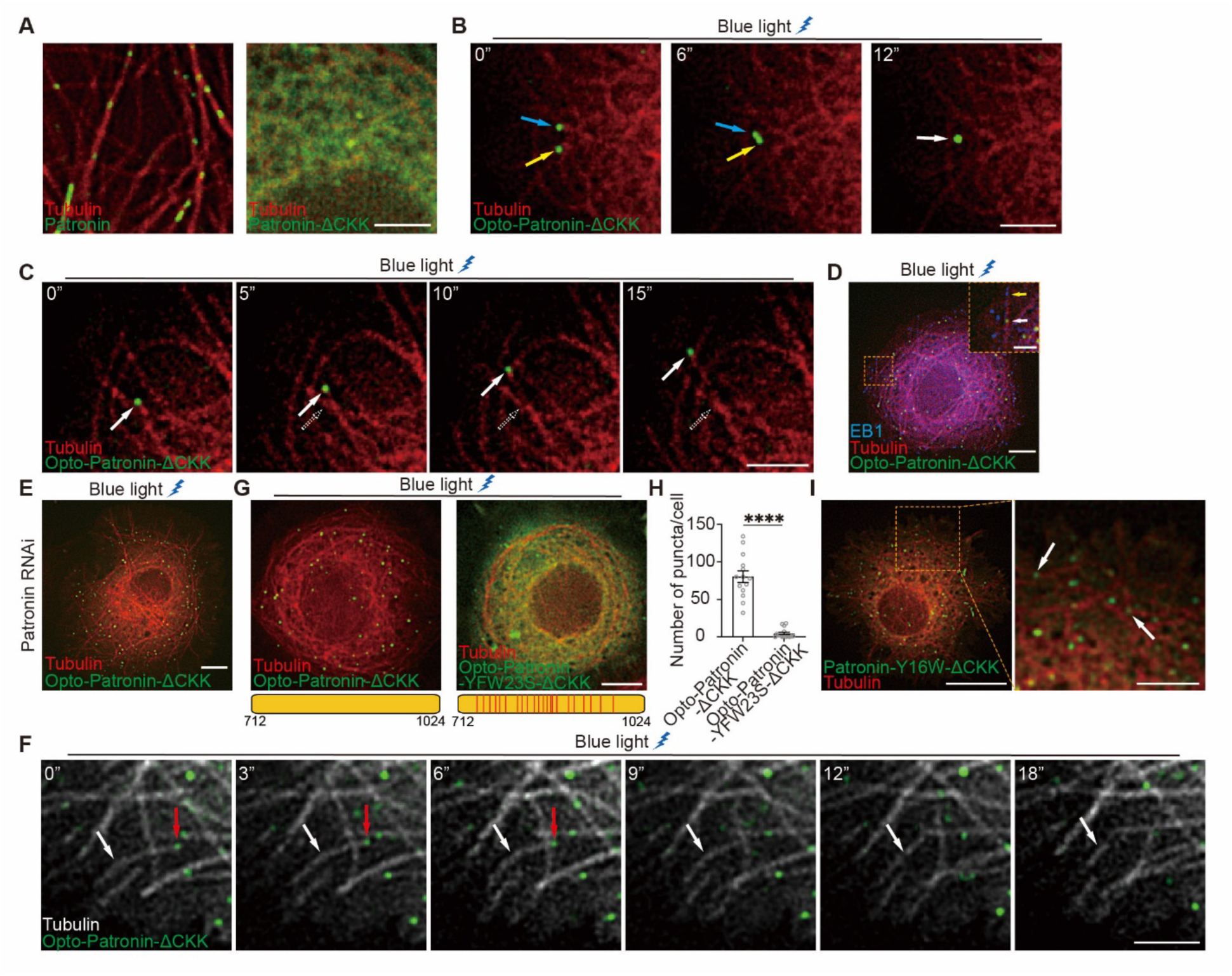
A second microtubule-recognition module cooperates with PrLD condensation to track minus ends. (A) Representative images of S2 cells expressing Patronin and Patronin-ΔCKK. Scale bar, 3 μm. (B) Live imaging of Opto-Patronin-ΔCKK condensates after blue light activation. Yellow and blue arrows, droplets before and during fusion; white arrows, droplet after fusion. Scale bar, 3 μm. (C) Live imaging of Opto-Patronin-ΔCKK condensates tracking microtubule minus ends after blue light activation. Solid arrows, Opto-Patronin-ΔCKK condensates at minus ends; dashed arrows, initial positions of the condensates. Scale bar, 3 μm. (D) Representative images of S2 cells co-expressing EB1 and Opto-Patronin-ΔCKK. Orange boxed region is magnified in the top right. Yellow arrows, EB1; white arrows, Opto-Patronin-ΔCKK condensates. Scale bars, 5 μm in main image and 2 μm in the magnified view. (E, F) Representative image (E) and live imaging (F) of Patronin-depleted S2 cells expressing Opto-Patronin-ΔCKK after blue light activation. White arrows, treadmilling microtubules; red arrows, Opto-Patronin-ΔCKK condensates tracking microtubule minus ends. Scale bars, 5 μm (E), 3 μm (F). (G, H) Representative images of S2 cells expressing Opto-Patronin-ΔCKK or Opto-Patronin-YFW23S-ΔCKK after blue light activation (G). Scale bar, 5 μm. (H) Quantification of number condensates. Data are presented as mean ± SEM; n = 14. *\*\*\*P < 0.001*. (I) Representative images of S2 cells expressing Patronin-Y16W-ΔCKK. White arrows, Patronin-Y16W-ΔCKK condensates at microtubule minus ends. Scale bars, 10 μm (left), 3 μm (right).

To test dynamic tracking, we used Patronin depletion to induce short treadmilling microtubules. Opto-Patronin-ΔCKK condensates remained associated with and tracked moving minus ends, but minus-end depolymerization continued and treadmilling was not suppressed (Fig. 6E, F). Thus, Opto-Patronin-ΔCKK can recognize and track minus ends but cannot efficiently stabilize or cap them. These data reveal a CKK-independent recognition activity while confirming an essential contribution of CKK to efficient minus-end stabilization. Truncation analysis identified residues 712-1457 as the minimal CKK-independent region supporting minus-end recognition (Extended Data Fig. 5B, C). This fragment contains the PrLD and a downstream region spanning residues 1025-1457. Because isolated PrLD does not bind microtubules, we hypothesized that the downstream region provides CKK-independent recognition information, whereas the PrLD supplies multivalent interactions that amplify it. Manipulating PrLD interaction strength supported this model. Aromatic-residue mutation strongly impaired condensation of Opto-Patronin-YFW23S-ΔCKK (Fig. 6G, H), whereas strengthening PrLD interactions by tyrosine-to-tryptophan substitution caused Patronin-Y16W-ΔCKK to condense spontaneously without optogenetic induction and selectively localize to minus ends (Fig. 6I). Thus, PrLD multivalency is required for stable minus-end enrichment by the non-CKK recognition module. Together, Patronin contains at least two separable sources of minus-end recognition information, CKK and a non-CKK region, neither of which efficiently produces robust targeting alone. PrLD-driven condensation amplifies these weak structural preferences into stable, persistent minus-end recognition and tracking without itself encoding microtubule polarity.

### Patronin condensates form a scaffold for cytoskeletal regulators at microtubule minus ends

Having established that Patronin condensation enables microtubule minus-end recognition, tracking and protection, we next asked whether Patronin condensates also function as scaffolds for additional cytoskeletal regulators. Patronin was previously reported to colocalize with the centrosomal protein Sas-4 in *Drosophila* S2 cells and was thus proposed to function as a scaffold at microtubule-organizing sites ^15^. Consistently, co-expressed Patronin and Sas-4 strongly colocalized (Extended Data Fig. 6A), whereas the condensation-defective Patronin-YFW23S mutant failed to efficiently recruit Sas-4 (Extended Data Fig. 6B). Replacing the Patronin PrLD with the unrelated phase-separating IDR from HNRNPA1 restored Sas-4 colocalization (Extended Data Fig. 6C), indicating that Sas-4 recruitment depends on Patronin condensation rather than the specific sequence of its PrLD. We next tested whether Patronin condensates recruit factors involved in microtubule remodeling. CAMSAP/Patronin family proteins interact with the microtubule-severing complex Katanin and contribute to the organization of non-centrosomal microtubule arrays ^17, 24, 42, 43^. Co-expression of Patronin with Katanin60 and Katanin80 resulted in strong enrichment of Katanin in Patronin condensates (Extended Data Fig. 6D). This recruitment was lost in Patronin-YFW23S (Extended Data Fig. 6E) but restored by Patronin–HNRNPA1 (Extended Data Fig. 6F). Thus, Patronin condensation provides a platform for Katanin recruitment. By concentrating a microtubule-severing factor together with a protein that captures and protects newly generated minus ends, Patronin condensates could spatially couple microtubule severing with minus-end stabilization.

We then asked whether this scaffolding function extends to the actin cytoskeleton. Recent work showed that Patronin and the Ena/VASP-family protein Ena cooperate in actin cable formation during *Drosophila* oogenesis ^44^. When Patronin and Ena were co-expressed in S2 cells, Ena condensates were recruited to Patronin assemblies associated with microtubules, forming prominent co-condensates that could fuse with one another (Fig. 7A, B). Ena recruitment was impaired by Patronin-YFW23S (Fig. 7C) and restored by Patronin–HNRNPA1 (Fig. 7D), again demonstrating a requirement for Patronin condensation. Ena/VASP proteins promote persistent actin barbed-end elongation and antagonize filament capping. Patronin-mediated recruitment of Ena could therefore locally promote the formation and extension of long actin filaments. Together, these findings suggest that Patronin condensation converts a microtubule minus-end-binding protein into a multifunctional cytoskeletal scaffold. By recruiting Katanin and Ena, Patronin condensates could provide a mechanism for spatially coupling minus-end stabilization and microtubule severing during microtubule remodeling, or for spatially coupling minus-end stabilization with actin assembly during microtubule-actin crosstalk.

**Fig. 7.**
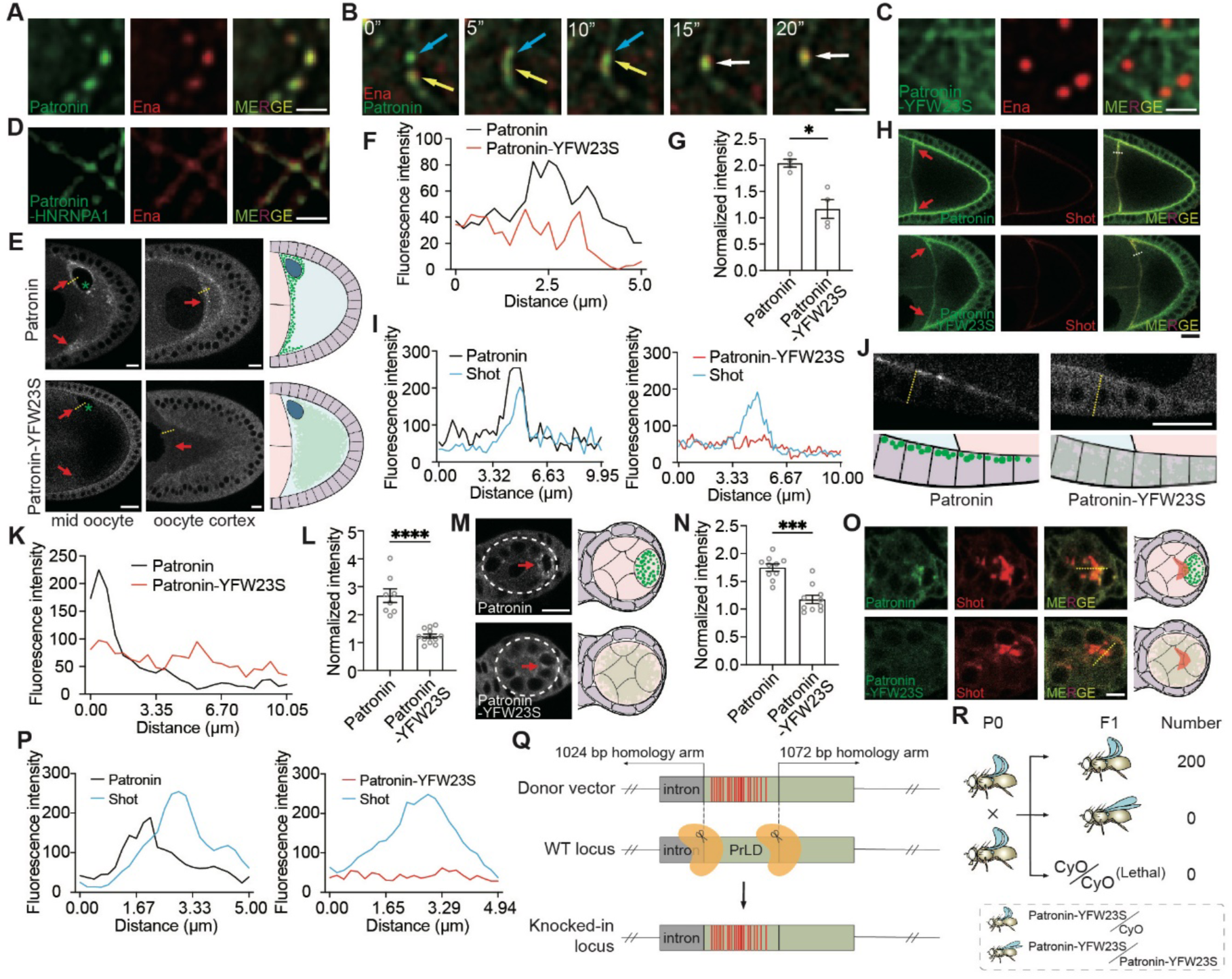
PrLD-mediated condensation is required for scaffolding, asymmetric localization and organismal development. (A, B) Representative images (A) and live imaging (B) of S2 cells co-expressing Ena and Patronin. Yellow and blue arrows, condensates before and during fusion; white arrows, condensates after fusion. Scale bar, 1 μm. (C, D) Disruption of phase separation in Patronin-YFW23S disrupts colocalization of Ena and Patronin, whereas introducing HNRNPA1 (Patronin-HNRNPA1) restores their colocalization. Scale bar, 1 μm. (E-G) Images of *ubi*-promoter driven Patronin and Patronin-YFW23S in stage 9 egg chambers at different focal planes (mid and cortex) of the oocyte (E). Red arrows point to the anterior region of the oocyte. Right, schematic of Patronin and Patronin-YFW23S distribution in the oocyte. Scale bar, 10 μm. (F) Patronin and Patronin-YFW23S fluorescence intensity along the yellow dashed line in the mid oocyte in (E). 0 at x-axis indicates the left end of the dashed line. (G) Quantification of Patronin and Patronin-YFW23S fluorescence intensity at mid oocyte focal plane, n = 4. (H, I) Representative images (H) and their quantification (I) of distribution patterns of *ubi*-promoter driven Patronin-mNG or Patronin-YFW23S-mNG together with endogenous Shot. Shot was labeled with an anti-Shot antibody. Scale bar, 20 μm. Fluorescence intensity in (I) was measured along the yellow dashed lines in (H). (J-L) (J) Distribution patterns of Patronin-mNG or Patronin-YFW23S-mNG in stage 9 follicle cells. Patronin-mNG is localized in the apical region of follicle cells. Bottom, schematic of Patronin and Patronin-YFW23S distribution in follicle cells. Scale bar, 20 μm. (K-L) Quantification of Patronin and Patronin-YFW23S fluorescence intensity along the yellow dashed line in (J). 0.00 at the x-axis (K) indicates the most apical position. (M, N) Distribution patterns (M) and Quantification (N) of Patronin-mNG or Patronin-YFW23S-mNG in region 3 of germarium, where germline cyst and follicle cells are initially derived. Right, schematic of Patronin and Patronin-YFW23S distribution in the region 3 of germarium. Scale bar, 10 μm. (O, P) Distribution patterns (O) and Quantification (P) of Patronin-mNG or Patronin-YFW23S-mNG together with Shot in region 3 of germarium. Right (O), schematic of Patronin and Patronin-YFW23S distribution with Shot. Scale bar, 5 μm. (Q) Schematic of CRISPR–Cas9-mediated genome engineering to generate endogenous Patronin PrLD mutations in *Drosophila*. Yellow, sgRNA–Cas9 complexes. (R) Schematic of Drosophila genetic experiments. Notably, the Patronin-YFW23S mutant is homozygous lethal.

### PrLD-mediated condensation is required for Patronin localization and developmental function *in vivo*

The results above establish that PrLD-mediated phase separation is required for Patronin to recognize, track and stabilize microtubule minus ends and to function as a molecular scaffold. We therefore asked whether this condensation activity is also required for Patronin localization and function *in vivo*. To address this question, we first generated transgenic flies expressing either Patronin-mNG or the condensation-deficient Patronin-YFW23S-mNG mutant under the control of the ubiquitin promoter. During stage 9 of *Drosophila* oogenesis, Patronin is recruited by Shot—the sole *Drosophila* spectraplakin capable of interacting with both F-actin and microtubules— to the anterior cortex of the oocyte, where microtubule minus ends are concentrated and non-centrosomal microtubule-organizing centers are assembled ^24^. Consistent with previous studies, Patronin-mNG was strongly enriched at the anterior oocyte cortex and displayed a punctate distribution. In contrast, Patronin-YFW23S-mNG failed to accumulate efficiently at the anterior cortex (Fig. 7E–G and Extended Data Fig. 6G, H). Immunostaining for Shot further showed that wild-type Patronin-mNG colocalized with Shot at the anterior cortex, whereas Patronin-YFW23S-mNG largely lost this colocalization (Fig. 7H, I). Importantly, Shot localization itself was comparable in oocytes expressing Patronin-mNG or Patronin-YFW23S-mNG, indicating that disruption of Patronin condensation impairs Patronin recruitment rather than the upstream localization of Shot. Thus, PrLD-mediated condensation is required for efficient Patronin enrichment at the Shot-defined anterior microtubule-organizing domain.

Patronin also localizes to the apical cortex of follicle cells during stage 9 oogenesis ^24, 25^. We found that apical enrichment of Patronin-YFW23S-mNG in follicle cells was strongly reduced compared with wild-type Patronin-mNG (Fig. 7J-L). Thus, the localization requirement for PrLD-mediated condensation is not restricted to the oocyte anterior but extends to another physiologically distinct cortical Patronin population. We next examined an earlier developmental context in which Patronin contributes to oocyte fate specification. During germline cyst development, Patronin becomes selectively enriched in the future oocyte, and this asymmetric localization has been proposed to depend on its ability to recognize and stabilize microtubule minus ends ^26^. Whereas wild-type Patronin-mNG was preferentially enriched in the future oocyte, Patronin-YFW23S-mNG showed markedly reduced enrichment and a more diffuse distribution (Fig. 7M, N). The condensation-deficient mutant also lost the prominent colocalization with Shot observed for wild-type Patronin-mNG in the presumptive oocyte (Fig. 7O, P). These observations indicate that PrLD-mediated condensation contributes to Patronin asymmetry during oocyte specification as well as to its later cortical localization.

The widespread localization defects of Patronin-YFW23S suggested that PrLD-mediated phase separation may be critical for a variety of Patronin-dependent cellular functions during development. To test this directly, we used CRISPR–Cas9 genome editing to replace all 23 aromatic residues within the Patronin PrLD with serine (Patronin-YFW23S; Fig. 7Q). Strikingly, homozygous Patronin-YFW23S mutant animals were lethal (Fig. 7R), with lethality occurring before the third larval instar. Thus, disrupting the multivalent interactions that drive Patronin condensation causes severe developmental defects. Together, these findings demonstrate that PrLD-mediated condensation is required for proper Patronin localization across multiple developmental contexts and is essential for normal organismal development. Combined with our aforementioned mechanistic analyses, these *in vivo* results support a model in which condensation is not merely a biochemical property of Patronin but a fundamental feature that enables its spatial organization and biological function.

## Discussion

Microtubule minus ends are essential determinants of microtubule organization, yet the molecular principles that specify, stabilize and organize these ends remain considerably less understood than those operating at plus ends. Here, we identify Patronin condensation as a mechanism that integrates these functions into a single, spatially restricted microtubule minus-end compartment. A prion-like domain (PrLD) provides the multivalent interactions that drive Patronin condensation but does not itself encode microtubule polarity. Instead, PrLD-mediated condensation cooperates with distinct microtubule-binding regions within Patronin to amplify weak or broadly distributed molecular preferences into robust minus-end recognition and dynamic tracking. The resulting Patronin-rich assembly functions as a protective cap that stabilizes minus ends against depolymerization and as a molecular scaffold that enriches additional cytoskeletal regulators, including Sas-4, Katanin and Ena. Importantly, disrupting PrLD-mediated condensation compromises Patronin localization in several developmental contexts and causes severe organismal defects. We therefore propose that Patronin condensation establishes a physiologically essential, multifunctional microtubule minus-end compartment that couples end recognition and tracking with stabilization, cytoskeletal organization and developmental function.

A central implication of our findings is that microtubule minus-end specificity does not reside in a single autonomous recognition domain. Structural studies of CAMSAP proteins have provided an attractive mechanism for CKK-mediated minus-end recognition, in which the CKK domain preferentially interacts with structural features associated with curved protofilaments near microtubule ends ^20^. However, this model has been difficult to reconcile with the behavior of isolated CKK domains in cells, where they frequently decorate the microtubule lattice rather than accumulating selectively at minus ends ^15, 22, 23^. Our results suggest that this discrepancy reflects a missing layer of regulation. CKK retains microtubule-binding and minus-end-recognition information, but in the cellular environment these interactions are insufficient to generate robust spatial asymmetry on their own. The PrLD provides the complementary and cooperating activity: it has little detectable affinity for microtubules by itself, but supplies strong multivalent interactions that cooperate with CKK. Linking PrLD or the heterologous HNRNPA1 to CKK is sufficient to reconstitute selective minus-end accumulation and dynamic tracking, whereas disrupting PrLD-mediated condensation abolishes this specificity. Thus, condensation acts as an amplifier of molecular recognition, transforming a modest structural preference into persistent spatial enrichment.

This principle may be conceptually related to mechanisms operating at microtubule plus ends. Plus-end identity is not specified by a single interaction but emerges from the cooperative behavior of networks of +TIPs that recognize nucleotide-and structure-dependent features of growing ends. Recent work has shown that multivalency within these networks can promote their collective accumulation at selected plus ends ^4–7^. A particularly instructive example is the budding-yeast Kar9 network, in which multiple interaction modules are distributed among three core +TIPs: Kar9, Bim1 (EB1) and Bik1 (Clip-170), whose redundant interactions generate a condensate-like +TIP body at specialized microtubule ends ^5^. Our results suggest that an analogous organizational principle may operate at the opposite end of the microtubule, but with much of the interaction network encoded within a single large, multidomain protein. Patronin contains distinct modules, including the CKK, PrLD and CC3 regions, that contribute different molecular properties; when multiple Patronin molecules assemble, these repeated modules can collectively generate a multivalent interaction network that promotes condensation and minus-end association. Thus, whereas multivalency in the Kar9 +TIP body is distributed across several interacting proteins, Patronin may integrate comparable modular complexity within repeated copies of one polypeptide to build a-TIP body, which is able to track the minus end ^16^. As a result, relatively weak molecular preferences can thereby be reinforced through cooperative interactions to generate an end-restricted molecular assembly. The underlying components and molecular cues differ, but the broader principle may be shared: microtubule-end identity can emerge from the combination of structural recognition and collective multivalent assembly (Fig. 8B), rather than from the stoichiometric one-to-one binding alone (Fig. 8A).

**Fig. 8.**
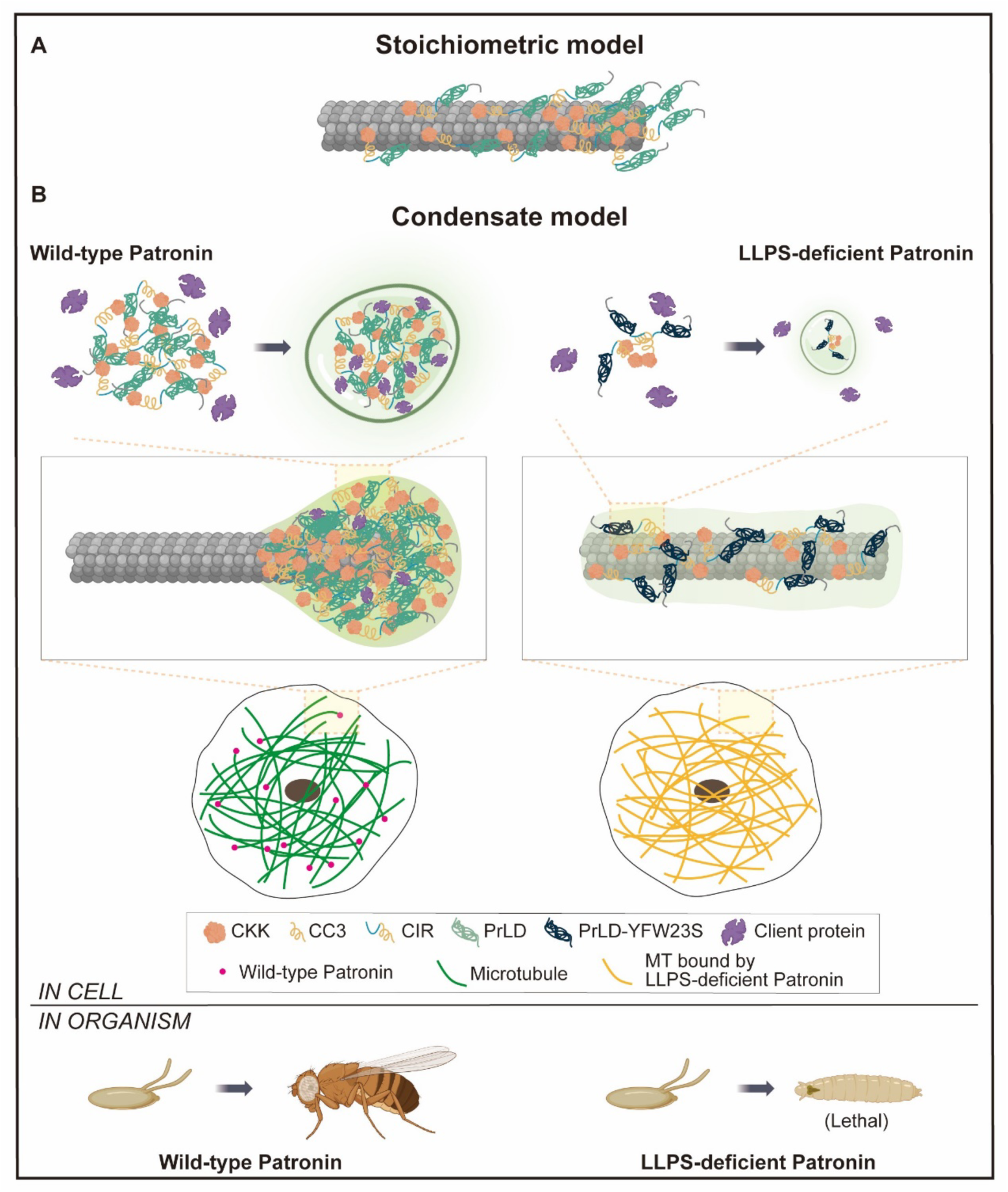
Condensation-based model for Patronin organization and function at microtubule minus ends. (A) Stoichiometric model. In the conventional model, Patronin associates with microtubule minus ends through discrete, stoichiometric interactions between individual Patronin molecules and the microtubule lattice/end structure. Patronin molecules are therefore envisioned primarily as individual minus-end-binding factors distributed along the terminal region of the microtubule. (B) Condensate model. Our results support an alternative model in which multivalent interactions among different Patronin domains, including the CKK, CC3, CIR and PrLD, promote collective assembly and PrLD-dependent condensation at microtubule minus ends. In wild-type Patronin, these interactions generate a cohesive Patronin-rich condensate that concentrates Patronin at the minus end and provides a multivalent platform for recruitment of client proteins such as Ena, Katanin, or Sas-4. Such condensation restricts Patronin to microtubule minus ends and supports its organization of the microtubule cytoskeleton in cells. In contrast, disruption of PrLD-mediated condensation, illustrated by the LLPS-deficient PrLD-YFW23S mutant, weakens collective Patronin assembly. LLPS-deficient Patronin consequently forms smaller assemblies (mediated by CC3) and redistributes more broadly along the microtubule lattice, despite retaining microtubule-binding activity (mediated by CKK and CIR). This altered spatial organization causes abnormal microtubule organization in cells and failure to support organismal development, resulting in early lethality. Together, the model proposes that Patronin function is not determined solely by stoichiometric binding to microtubule minus ends, but emerges from the combination of microtubule recognition and cooperative multivalent condensation, which spatially concentrates Patronin and associated client proteins at microtubule minus ends. Symbols: CKK, C-terminal kinesin-like calponin-homology domain; CC3, coiled-coil 3 region; CIR, CKK-independent microtubule recognition region; PrLD, prion-like domain; PrLD-YFW23S, LLPS-deficient PrLD mutant. Purple symbols indicate Patronin-associated client proteins. Green and yellow filaments represent microtubules organized by wild-type and LLPS-deficient Patronin, respectively.

The identification of a second, CKK-independent minus-end recognition activity further supports this view. Previous biochemical studies indicated that a central region of *Drosophila* Patronin can bind microtubule minus ends independently of CKK *in vitro* ^16^, but how this activity contributes to minus-end targeting in cells has remained unclear. We find that a central Patronin region containing the PrLD and a downstream microtubule-binding region can recognize and dynamically track minus ends when sufficient condensation is provided. Patronin lacking CKK can therefore retain minus-end recognition and tracking while losing efficient minus-end stabilization. These observations separate two properties that are often treated as equivalent: minus-end tracking and stabilization, and suggest that Patronin contains partially redundant recognition mechanisms that converge on a common condensation-dependent targeting process. Such redundancy may be advantageous because microtubule minus ends are unlikely to represent a single invariant structural state. Minus ends can differ in protofilament configuration ^20, 45, 46^, nucleotide state ^47, 48^, association with nucleation complexes ^49^, and history of severing or depolymerization ^15, 17^. Multiple recognition modules coupled through condensation could allow Patronin to remain associated with structurally heterogeneous or dynamically changing minus ends.

Our results also reveal that not all Patronin condensation is functionally equivalent. In addition to the PrLD, the conserved CC3 region supports weaker condensation. CC3-mediated interactions increase CKK association with microtubules and improve its ability to stabilize microtubules at low protein concentrations, but do not confer robust minus-end specificity. By contrast, PrLD-mediated condensation produces strong selective enrichment at minus ends. These findings suggest that the biological output of condensation depends not simply on whether a protein forms condensates, but on the strength and balance of protein-protein and protein-surface interactions. This distinction is evident in the different wetting behaviors of Patronin variants. Weakening PrLD-mediated cohesion favors extensive wetting or coating of the microtubule lattice, whereas strengthening aromatic interactions shifts Patronin toward more compact bead-like assemblies and reduced molecular mobility. Such behavior is consistent with general physical principles governing condensate wetting on biological surfaces ^50^. We therefore favor a model in which PrLD-mediated cohesion is in accordance with an intermediate state that permits Patronin to remain dynamically associated with the microtubule while resisting complete spreading along the lattice.

This framework also provides a physical interpretation of the long-standing concept of Patronin as a microtubule minus-end “cap.” Goodwin and Vale originally showed that Patronin protects minus ends from the Kinesin-13 depolymerase Klp10A and proposed that Patronin caps the minus end ^15^. The molecular architecture of such a cap, however, was not defined. A simple interpretation would be a small stoichiometric complex that occupies the terminal tubulin subunits. Our findings suggest a broader view. Patronin can form a concentrated assembly centered at the minus end and extending for a finite distance along the adjacent lattice, reminiscent of the CAMSAP “stretch”, which was previously shown for CAMSAP2 and CAMSAP3 ^17^. We propose that this assembly functions as a condensate-based protective cap. By concentrating multiple Patronin molecules around the end, condensation would increase the effective occupancy of microtubule-binding sites and create a locally protected zone that is resistant to depolymerizing activity. The functional data support this interpretation. PrLD-CKK is sufficient to form a minus-end-enriched assembly and suppress the treadmilling phenotype caused by Patronin depletion, whereas the condensation-defective PrLD-YFW23S-CKK fails to do so. CKK alone can stabilize microtubules when expressed at sufficiently high levels, indicating that CKK contains the core protective activity but that condensation strongly increases its effectiveness. Consistent with this view, CC3-mediated condensation increases CKK occupancy on microtubules and enhances stabilization even though it does not generate selective minus-end enrichment. PrLD-mediated condensation therefore contributes an additional level of spatial focusing by concentrating protective CKK activity specifically at the minus end. We do not exclude direct steric competition between CKK and Kinesin-13, as proposed from structural considerations ^20^. Rather, condensation could amplify such molecular competition by concentrating many CKK-containing Patronin molecules within the minus-end region. In this sense, the Patronin “cap” may be an emergent property of a mesoscale assembly rather than a single capping complex.

A condensate-based cap also provides a natural explanation for another poorly understood property of Patronin: its proposed scaffolding function. Goodwin and Vale observed that Patronin-containing structures associate with centrosome-related proteins including Sas-4 and suggested that Patronin might organize additional factors at sites of microtubule formation ^15^. Phase separation offers a straightforward physical mechanism for such organization because condensates can selectively enrich client molecules while maintaining dynamic exchange ^28^. Our data show that Patronin condensates enrich not only Sas-4 but also the microtubule regulator Katanin and the actin regulator Ena, and that this enrichment depends on intact PrLD-mediated condensation. Patronin condensates may therefore constitute local cytoskeletal organizing hubs rather than simply protective structures at filament ends. The association with Katanin and Ena suggests two potentially important outputs of this scaffolding activity. Microtubule severing generates new ends that must subsequently be stabilized or incorporated into the network; concentrating Katanin together with a minus-end protector could therefore spatially couple microtubule severing with capture and stabilization of newly generated minus ends. Ena/VASP proteins, in turn, promote persistent actin barbed-end elongation and actin network organization. The enrichment of Ena in Patronin condensates raises the possibility that minus-end-associated assemblies also provide sites where microtubule and actin regulation intersect. These models remain to be tested directly, but they suggest that the functional composition of Patronin condensates could be adapted to different cellular contexts, allowing a common condensation mechanism to coordinate distinct forms of cytoskeletal remodeling.

Our *in vivo* localization data add functional understanding of Patronin beyond *in vitro* systems and cultured cells. During stage 9 oogenesis, wild-type Patronin is strongly enriched at the Shot-defined anterior cortex of the oocyte, whereas Patronin-YFW23S fails to accumulate efficiently at this site. Importantly, Shot itself remains normally localized when Patronin condensation is disrupted. These observations are consistent with a model in which an upstream cortical landmark such as Shot provides spatial information or initial recruitment, while PrLD-mediated condensation amplifies and stabilizes Patronin accumulation into a persistent cortical domain. In this view, condensation does not determine where Patronin should localize; rather, it converts a local recruitment cue into robust mesoscale organization. The reduced apical localization of Patronin-YFW23S in follicle cells and its diminished enrichment in the future oocyte suggest that this principle operates across multiple developmental contexts rather than at a single specialized site. The developmental consequences of disrupting PrLD multivalency further underscore its physiological importance. Introducing the YFW23S substitutions at the endogenous Patronin locus causes homozygous lethality before the third larval instar, demonstrating that the aromatic interactions are required for normal development. It should be noted that these substitutions could in principle perturb specific molecular interactions in addition to condensation. However, our data argue that loss of multivalent condensation is the major underlying defect. First, PrLD deletion and aromatic-residue mutation produce similar cellular phenotypes in S2 cells. Second, strengthening PrLD interactions produces the opposite material-state shift. Third, a heterologous phase-separating HNRNPA1 IDR can restore Patronin’s minus-end tracking, stabilization and scaffolding activities in S2 cells.

Our findings also raise the question of how condensation has diversified within the CAMSAP/Patronin family. While this manuscript was in preparation, Hsieh et al. (2026) reported that the only Patronin/CAMSAP homolog in *C. elegans*, PTRN-1, forms dynamic condensates in intestinal epithelial cells in response to pore-forming toxin challenge ^33, 51^. PTRN-1 condensation is promoted by a TTC38–HSP70–HSP90 pathway and contributes to microtubule-dependent trafficking and epithelial recovery, providing independent *in vivo* evidence that condensation can have physiological functions in an invertebrate Patronin/CAMSAP family protein. However, it is not known which region or domain is required for PTRN-1 condensation, neither is it known whether PTRN-1 condensates bind microtubule minus-ends. Notably, our sequence analysis indicates that PTRN-1 and *Drosophila* Patronin share a similar architecture in which a PrLD is positioned upstream of CC3 and CKK (Extended Data Fig. 2A), whereas the three mammalian CAMSAPs retain CC3 and CKK but lack a comparable PrLD ^17, 18, 22^. Together with our finding that the Patronin PrLD provides a major driving force for condensation whereas CC3 makes a weaker contribution, this suggests that the molecular basis of condensation may have diversified during vertebrate CAMSAP expansion. Mammalian CAMSAP2 undergoes phase separation *in vitro*, with its CC3–CKK region contributing to condensate-associated, γ-TuRC-independent microtubule nucleation ^32^, indicating that a PrLD is not essential for this property. The expansion of the vertebrate family to CAMSAP1–3 (compared to only one homolog in *C. elegans* and *Drosophila*) raises the possibility that interactions among CAMSAP paralogs and additional partners provide network-level multivalency that reinforces weaker condensation activities encoded by individual proteins. Indeed, CAMSAP2 and CAMSAP3 can cocluster and cooperate at noncentrosomal microtubule minus ends ^52^, although whether such interactions promote condensate formation remains unknown. In this sense, invertebrate and vertebrate CAMSAPs may represent different implementations of a common organizational principle: multivalent interactions can be either concentrated within repeated copies of a single PrLD-containing protein or distributed across a network of cooperating proteins, analogous to the multicomponent Kar9–Bim1–Bik1 +TIP body. Determining whether mammalian CAMSAPs form such collective mesoscale assemblies at endogenous concentrations, and how these assemblies regulate minus-end targeting, protection and client recruitment, will be important questions for future work.

Several mechanistic questions remain open. First, the structural cue recognized by Patronin’s CKK-independent microtubule-binding region is unknown, as is its relationship to the structural preference of CKK. Second, determining whether and how Katanin, Ena, Sas-4 and other factors interact directly with Patronin or are recruited will be important for defining the scaffold function of the compartment. Finally, the *in vivo* data raise the question of how developmental landmarks such as Shot cooperate with microtubule-end cues and Patronin condensation to set the size, position and material properties of endogenous Patronin assemblies. Together, our findings suggest a revised view of microtubule minus-end organization. Patronin does not simply recognize a terminal tubulin structure and cap it through a stoichiometric binding event. Instead, weak minus-end preferences encoded by multiple microtubule-binding regions are amplified by PrLD-driven multivalent condensation to generate a dynamic, spatially restricted Patronin compartment. This compartment provides persistent minus-end recognition and tracking, functions as a protective cap against depolymerization, creates a scaffold for additional cytoskeletal regulators and, *in vivo*, enables Patronin to assemble at developmentally specified sites. The requirement for PrLD multivalency for normal Patronin localization and organismal viability further indicates that condensate-based compartmentalization is not merely a biochemical property but a physiologically essential mechanism for establishing stable and multifunctional microtubule minus-end identity.

## Acknowledgements

We are grateful to Dr. Vladimir Gelfand, Dr. Daniel St Johnston, Bloomington Drosophila Stock Center for fly stocks. We want to thank Dr. Wenyu Wen for her helpful discussion of the manuscript. This work is supported by grant from the National Natural Science Foundation of China (32671108) to JC.

## Author contributions

Conceptualization: JC, BZ, HW; Methodology: BZ, HW, KZ, JY, SY, JC, XL, LQH, LH; Investigation: BZ, JY, SY, HW, KZ; Resources: BZ, HW, JY, KZ, JC, SY, XL, LQH, LH; Visualization: BZ, SY, HW; Writing and Editing: JC, BZ, HW, XL, LQH, LH; Supervision: JC and HW.

## Methods

### S2 cell culture and transfection

*Drosophila* S2 cells were cultured at 25 °C in Schneider’s Drosophila medium (Gibco, 21720024) supplemented with 10% fetal bovine serum (FBS; Gibco, A5670701) and 1% penicillin–streptomycin. Plasmids were transfected using Effectene (Qiagen, 301425) according to the manufacturer’s instructions.

For Patronin RNAi, dsRNA targeting the Patronin 5′ UTR was synthesized using the T7 RNAi Transcription Kit (Vazyme, TR102), with S2 cell cDNA as the template and the following primers:

Forward, 5′-TAATACGACTCACTATAGGGCACATGAAAATTTGTAAG-3′; Reverse, 5′-TAATACGACTCACTATAGGGGACTCCGGCTCTCCGACGCCCGCC-3′ ^16^. S2 cells (0.5 × 10⁶ per well) were plated in 24-well plates and transfected the next day with dsRNA, pMT and pAC plasmids using Effectene. Expression from pMT vectors was induced with 25-200 μM CuSO₄ for 18-24 h.

### Plasmid construction

*Drosophila* codon-optimized mNeonGreen ^53^ was inserted into a metallothionein promoter vector to generate pMT-mNG. Patronin, amplified from a *Drosophila* cDNA pool, or its mutant variants were inserted downstream of mNG. For microtubule imaging in S2 cells, mCherry-αTub84B was cloned into pAC5.1. For ubiquitous expression in flies, the UAS cassette of pUASTattB (Addgene, 36431) was replaced with the *Drosophila* Ubiquitin p63E promoter to generate pUbi-attB ^54^. Patronin– mNG or mutant Patronin–mNG sequences were inserted downstream of this promoter.

Patronin truncation–EGFP constructs were cloned into pET28a using the ClonExpress Ultra One Step Cloning Kit V2 (Vazyme, C116) for expression in *Escherichia coli* BL21(DE3). For expression in Sf9 cells, PCR-amplified MBP– Patronin–EGFP–His or corresponding mutant sequences were inserted into pFastBacHT C using RsrII and BamHI. The sequence encoding human hnRNPA1 residues 186–319 was amplified from human cDNA. Two tandem copies of this sequence replaced the Patronin prion-like domain in mNG–Patronin or MBP– Patronin–GFP–His constructs.

### Patronin expression and purification in insect cells

Patronin proteins were expressed using the baculovirus system. pFastBacHT C constructs were transformed into DH10Bac *E. coli* (Tsingke Biotechnology, DLC104), and recombinant bacmids were isolated after blue–white screening and verified by PCR. Sf9 cells were maintained at 27 °C in SIM SF medium (Sino Biological, MSF1).

Sf9 cells (1.0 × 10⁶ cells ml⁻¹) were transfected with recombinant bacmids using Lipofectamine 3000 (Invitrogen, L3000015), and P1 virus was harvested after 72 h. Virus was amplified by infecting 15 ml of Sf9 cells with P1; P2 virus was harvested 4 days after proliferation arrest. Typically, 100 ml of Sf9 cells (1.5 × 10⁶ cells ml⁻¹) were infected with P2 at a 1:10 ratio and harvested after 48 h. Cells were washed, resuspended in harvest buffer (25 mM HEPES, pH 7.6, 150 mM KCl and 1× EDTA-free protease inhibitor cocktail; MCE, HY-K0011), snap-frozen in liquid nitrogen and stored at-80 °C. All subsequent steps were performed at 4 °C. Cells were disrupted with sonication. Sf9 cells expressing recombinant proteins were resuspended in lysis buffer and sonicated. The base purification buffer contained 25 mM HEPES, pH 7.6, 500 mM KCl, 0.1% CHAPS, 10% glycerol, 0.5 mM DTT and 2 mM MgCl_₂_; lysis buffer additionally contained 5 mM imidazole and 1× protease inhibitor cocktail. Lysates were clarified at 40,000 × g for 30 min, and proteins were captured on Ni– NTA resin (Smart-Lifesciences, SA005025). Resin was washed five times with base buffer containing 10 mM imidazole, and proteins were eluted with base buffer containing 500 mM imidazole. Proteins were dialysed overnight against 25 mM HEPES, pH 7.6, 500 mM KCl, 10% glycerol, 1 mM DTT and 2 mM MgCl_₂_, and analyzed by SDS–PAGE.

### Patronin truncation expression and purification in bacteria

Patronin truncations were expressed in E. coli BL21(DE3) by induction with 0.2 mM IPTG for 16 h at 18 °C. Cell pellets were snap-frozen in liquid nitrogen, resuspended in lysis buffer (25 mM HEPES, pH 7.6, 300 mM KCl, 5 mM imidazole, 10% glycerol and 2 mM MgCl_₂_) supplemented with protease inhibitors, and sonicated. Lysates were clarified at 12,000 × g for 40 min at 4 °C, and proteins were purified on Ni-NTA resin as described above.

### *In vitro* phase separation assays

Phase separation assays were performed in 25 mM HEPES, pH 7.6, 150 mM KCl, 2 mM MgCl_₂_ and 1 mM DTT without crowding agents unless specified in the figure legends. Protein concentrations were adjusted to yield a final KCl concentration of 150 mM. Frozen protein aliquots were thawed and centrifuged at 13,000 rpm for 5 min to remove aggregates. For Patronin-GFP and its mutants, MBP tags were cleaved during the phase separation reaction by incubation overnight at 4 °C with 1 U PreScission protease (Beyotime, P2302) per 30 μl reaction. Phase separation assay was performed in Eppendorf tubes, and content from tubes were immediately transferred to glass-bottom dishes (NEST Biotechnology). After up to 10 min to allow droplets to settle, samples were imaged on a Leica SP5 confocal microscope. Condensates were counted in 50 × 50 μm regions using ImageJ.

### *Drosophila* stocks and genetics

*Drosophila* melanogaster were maintained on standard medium at 25 °C and 60% relative humidity. *UASp-GFP-Patronin* was provided by Vladimir Gelfand (Northwestern University, USA)^34^, and *Patronin-YFP* by Daniel St Johnston (University of Cambridge, UK)^24^. *Mat-Gal4* (Bloomington Drosophila Stock Center, 7063) drove germline expression of *UASp-GFP-Patronin*. *Ubi-Patronin-mNG* and *Ubi-Patronin-YFW23S-mNG* transgenic lines were generated by phiC31-mediated site-specific integration at attP2.

The *Patronin-YFW23S* knock-in strain was generated by injecting *w¹¹¹⁸* embryos with *in vitro*-transcribed sgRNAs (Promega, P1320), a donor plasmid and Cas9 protein (NEB, M0386). Four sgRNAs were used:

sgRNA1, 5′-GGTCTGCTGCTGTGGATGCA-3′;

sgRNA2, 5′-GATGCACGGTTAGTGTGGGC-3′;

sgRNA3, 5′-CCTCGCCACAGCATGGCCAA-3′;

sgRNA4, 5′-TGGATCCACTAACCTGCCGT-3′. The 30 μl injection mixture contained sgRNA at 150 ng μl⁻¹, Cas9 at 500 ng μl⁻¹ and donor plasmid DNA at 500 ng μl⁻¹.

### Live imaging of S2 cells and egg chambers

CuSO₄-induced S2 cells were plated on glass-bottom dishes (NEST Biotechnology) coated with 0.5 mg ml⁻¹ concanavalin A (ConA; Sigma-Aldrich, C7275) in water and air-dried. Cells were incubated at 25 °C for 1 h to adhere and spread. Live imaging of S2 cells on coverslips was performed on a GE DeltaVision Elite system with a 100×/1.4 NA oil-immersion objective at 5 s intervals. Images were deconvolved using DeltaVision Elite software and processed in ImageJ. For two-colour images, channel brightness was adjusted separately before merging.

Egg chambers from well-fed, 3–5-day-old females were dissected and isolated in Schneider’s medium (Gibco, 21720024) supplemented with 15% FBS (Gibco, A5670701) and 0.1 mg ml⁻¹ insulin, as previously described ^55^. Isolated egg chambers were transferred to eight-well chambered coverglass (Thermo Fisher Scientific, 155411) precoated with 0.2 mg ml⁻¹ poly-D-lysine (Beyotime, ST508) to prevent movement and imaged in the same medium ^56^. Images were acquired at 20 s intervals on a ZEISS LSM880 confocal microscope with a 40×/1.3 NA oil-immersion objective and processed using Zeiss ZEN and ImageJ.

### Immunofluorescence staining

Ovaries were dissected at room temperature in PBS, pH 7.4, and egg chambers were fixed for 10 min in 4% paraformaldehyde, pH 7.4 (Beyotime, P0099). Samples were washed three times in PBT (PBS containing 0.1% Triton X-100; Sangon, A110694). For F-actin labelling, samples were incubated with TRITC-phalloidin (1:500; Yeasen, 40734ES75) for 60 min and washed three times in PBT. For antibody staining, samples were blocked for 30 min in PBT containing 10% goat serum (Boster, AR1009), incubated with primary antibodies overnight at 4 °C, washed three times in PBT, and incubated with secondary antibodies (1:200, Jackson ImmunoResearch) for 2 h at room temperature. Primary antibody was as followed: mouse monoclonal anti-Shot antibody (shot mAbRod1, DSHB, 1:5). After three further PBT washes, samples were imaged on a ZEISS LSM880 confocal microscope.

### Fluorescence recovery after photobleaching

Fluorescence recovery after photobleaching (FRAP) was measured in egg chambers, S2 cells and in vitro condensates on a Leica SP5 confocal microscope with a 63×/1.4 NA oil-immersion objective. Patronin and its mutants were bleached with a 488 nm laser, and microtubules in S2 cells with a 561 nm laser. After one prebleach image, a circular region of interest encompassing the condensates was bleached at 100% laser power. Images were acquired at 2 s intervals for 60 s. Fluorescence in the bleached region was normalized to its prebleach intensity (100%) and plotted using GraphPad Prism.

### Optogenetic droplet induction

The optoDroplet assay was adapted from a published protocol ^57^. Transfected S2 cells were plated on ConA-coated glass-bottom dishes as described above and cultured in the dark. Images were acquired on a GE DeltaVision Elite system with a 100×/1.4 NA oil-immersion objective. Blue light (461-480 nm) induced droplet formation during imaging. Images were deconvolved using DeltaVision Elite software and processed in ImageJ.

## Statistical analysis

Data were analysed using GraphPad Prism 10. Two-group comparisons used unpaired, two-tailed Student’s t-tests, and multiple-group comparisons used one-way ANOVA. Data are presented as mean ± s.e.m. Significance thresholds were *\*P < 0.05, **P < 0.01, ***P < 0.001 and ****P < 0.0001*.

