## Supplementary data for "Patronin condensation integrates recognition, protection and organization of microtubule minus ends"

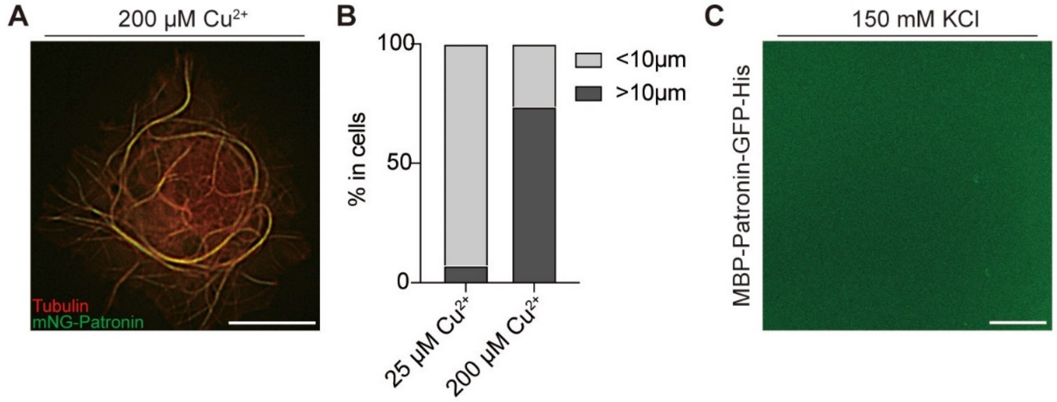


**Extended Data Fig. 1. High levels of Patronin causes spreading of Patronin condensate along microtubule lattice**

(A) Patterns of mNG-Patronin on microtubules in S2 cells induced with 200 μM $Cu^{2+}$. Scale bar, 10 μm.

(B) Quantification of Patronin condensate length along microtubules under induction with 25 μM and 200 μM $Cu^{2+}$. n=42.

(C) Representative image of 3 μM MBP-Patronin-GFP-His under 150 mM KCl conditions in vitro, indicating that MBP-Patronin-GFP purified from Sf9 cells does not form condensate before MBP removal. Scale bar, 10 μm.

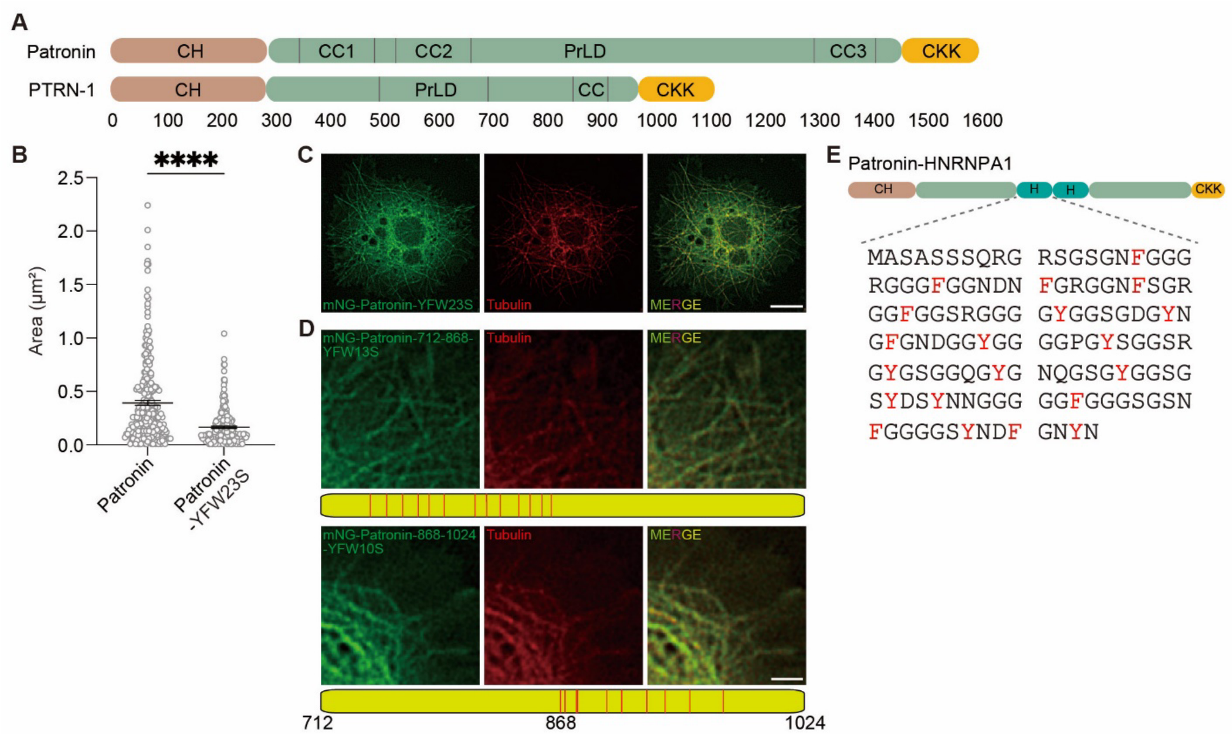


**Extended Data Fig. 2. Schematic of domain arrangement of Patronin, PTRN-1 and Patronin-HNRNPA1, and reduced condensation by mutations within PrLD**

(A) The *C. elegans* PTRN-1 is very similar to *Drosophila* Patronin in domain arrangement.
(B) Comparison of droplet areas between Patronin-GFP (n = 294) and Patronin-YFW23S-GFP (n = 210) at 1 µM in 100 mM KCl. Data are mean ± s.e.m.
(C) Representative images of S2 cells expressing mNG-Patronin-YFW23S. Scale bar, 5 µm.
(D) Representative images of S2 cells expressing Patronin PrLD mutants with varying numbers of aromatic residues. Below the fluorescence images, schematics of the PrLD mutations are shown. Scale bar, 2 µm.
(E) The domain arrangement of the chimeric Patronin-HNRNPA1. The central PrLD domain is replaced by two tandem copies of the HNRNPA1's IDR region (abbreviated as H). The Amino acid sequence of one H is listed below, aromatic residues are highlighted in red. Note that 2 tandem copies of H contain a total of 268 amino acids and are comparable in size to that of PrLD (312 amino acids).

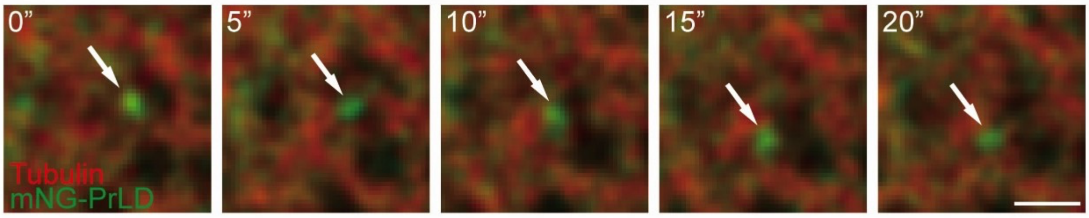


**Extended Data Fig. 3. Isolated PrLD does not associate with microtubule**
Time-lapse imaging of S2 cells expressing mNG-PrLD. White arrows, PrLD foci. Notably, PrLD foci did not bind to microtubules or track microtubule minus ends. Scale bar, 1 µm.

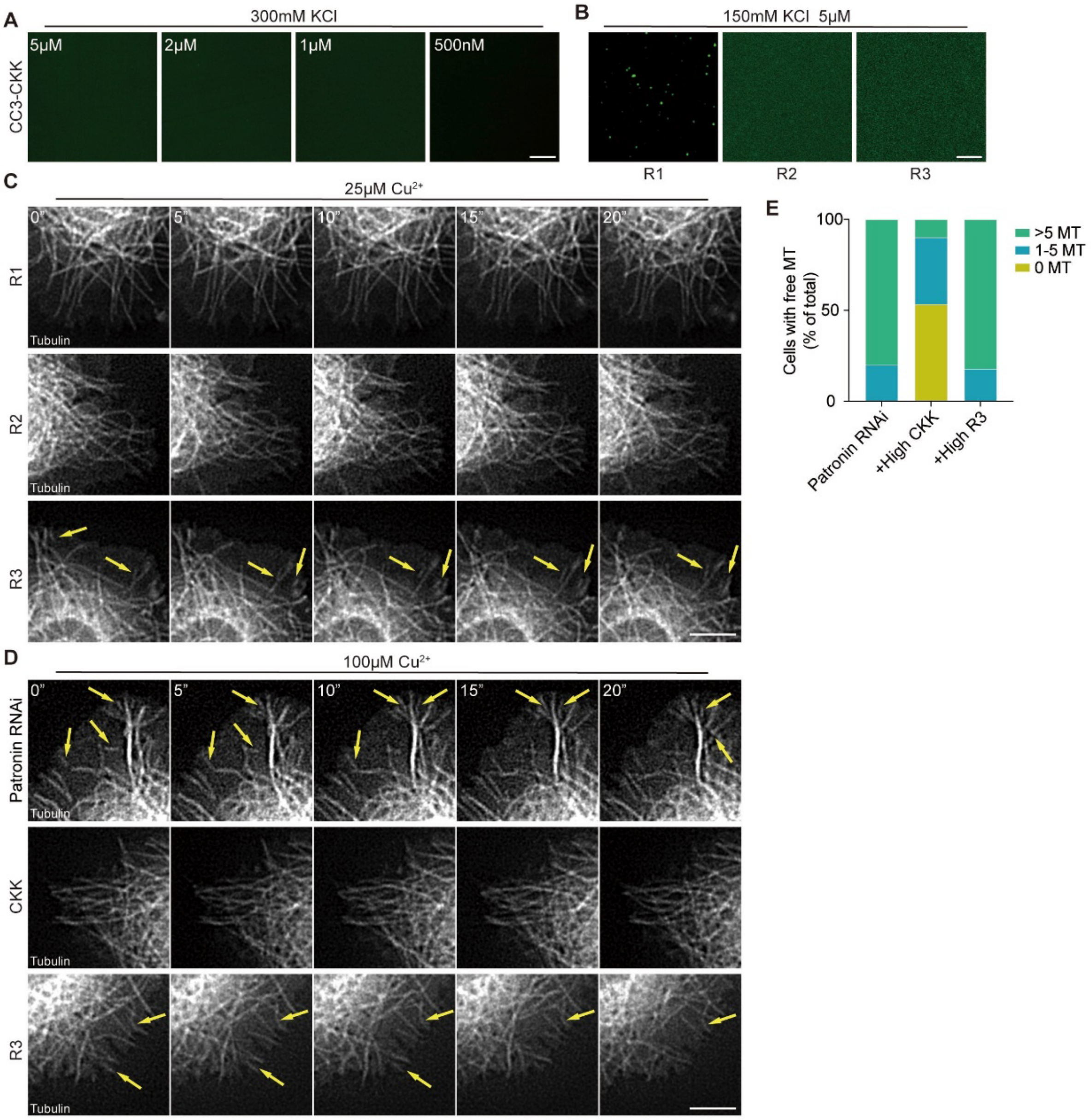


**Extended Data Fig. 4. Various truncations of CC3-CKK displayed different condensation abilities and rescue effects on treadmilling microtubules**

(A) Representative fluorescence images of CC3-CKK-GFP displaying no LLPS at different protein concentrations in an *in vitro* buffer containing 300 mM KCl. Scale bar, 10 μm.

(B) Representative *in vitro* LLPS images of CC3-CKK truncations without crowding agent. Scale bar, 5 μm.

(C) Time-lapse imaging of Patronin-depleted S2 cells expressing R1, R2 and R3 truncation constructs under induction with 25 μM $Cu^{2+}$; quantification is shown in Fig. 5L. Yellow arrows indicate “free microtubules.” Scale bar, 5 μm.

(D, E) Time-lapse microscopy of S2 cells expressing *Patronin RNAi* together with R3 or CKK under induction with 100 μM $Cu^{2+}$. Yellow arrows indicate “free microtubules.” Quantification is shown in (E). Scale bar, 5 μm.

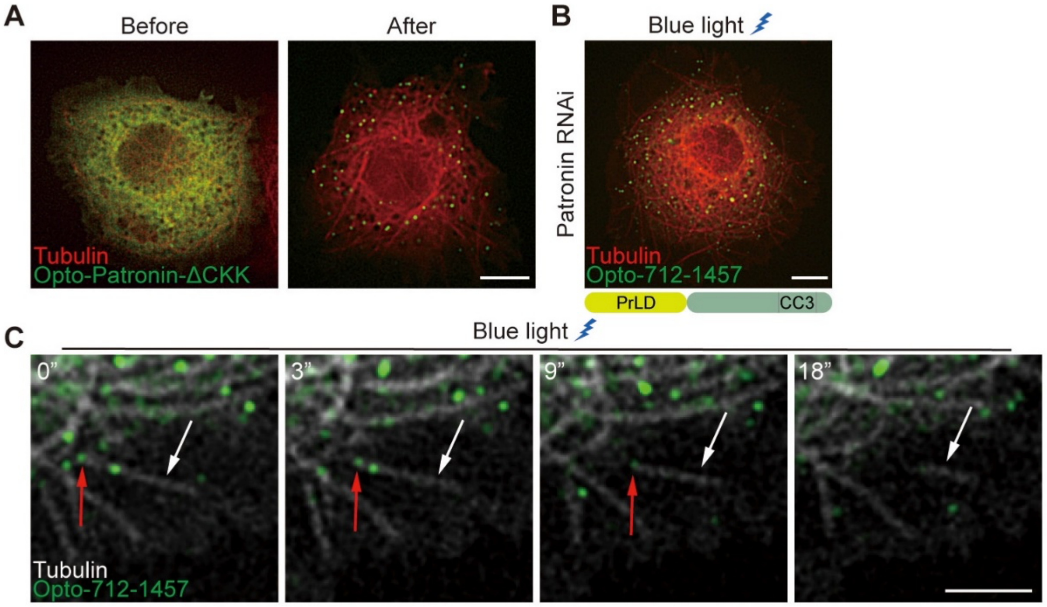


**Extended Data Fig. 5. CKK-independent recognition region tracked but failed to stabilize minus ends**

(A) Representative images of S2 cells expressing Opto-Patronin-ΔCKK before (left) and after (right) 30 s of blue-light illumination. Scale bar, 5 μm.

(B, C) Representative images (B) and live imaging (C) of Patronin-depleted S2 cells expressing Opto-712-1457 after 30 s of blue-light illumination. Bottom, schematic of residues 712–1457. Scale bar, 5 μm. White arrows, treadmilling microtubules; red arrows, Opto-712-1457 condensates track microtubule minus ends. Scale bar, 3 μm.

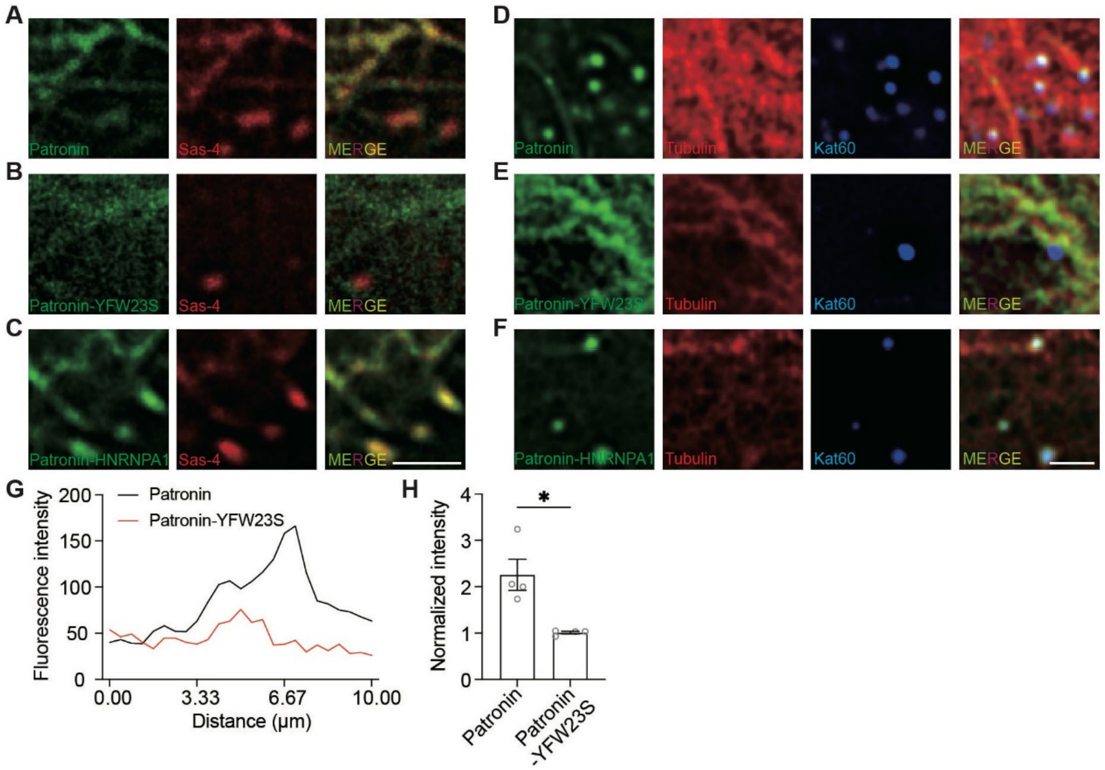


**Extended Data Fig. 6. Condensation of Patronin is required for colocalization of Sas-4, Katanin with Patronin**

(A-C) Representative images of S2 cells expressing Patronin, or Patronin-YFW23S, or Patronin-HNRNPA1 together with Sas-4. Notably, Sas-4 and Patronin-YFW23S do not co-localize. Scale bar, 2 μm.

(D-F) Representative images of S2 cells expressing Patronin, or Patronin-YFW23S, or Patronin-HNRNPA1 together with Kat60. Notably, Kat60 and Patronin-YFW23S do not co-localize. Scale bar, 1 μm.

(G, H) Quantification of Patronin and Patronin-YFW23S fluorescence intensity along the yellow dashed line in the oocyte cortex shown in Fig. 7E.
